# Stable kinetochore-microtubule attachment requires loop-dependent Ndc80-Ndc80 binding

**DOI:** 10.1101/2022.08.25.505310

**Authors:** Soumitra Polley, Helen Müschenborn, Melina Terbeck, Anna De Antoni, Ingrid R. Vetter, Marileen Dogterom, Andrea Musacchio, Vladimir A. Volkov, Pim J. Huis in ‘t Veld

**Author notes:** Centre for Medical Biotechnology, University of Duisburg-Essen, Essen, Germany (Helen Müschenborn). IFOM-The AIRC Institute of Molecular Oncology, Milan, Italy.

## Abstract

During cell division, kinetochores link chromosomes to spindle microtubules. The Ndc80 complex, a crucial microtubule binder, populates each kinetochore with dozens of copies. Whether adjacent Ndc80 complexes cooperate to promote microtubule binding remains unclear. Here we demonstrate that the Ndc80 loop, a short sequence identified across eukaryotes and predicted to interrupt the Ndc80 coiled-coil, promotes direct interactions between full-length Ndc80 complexes. Both in dividing cells and in a fully reconstituted system, these interactions are essential for the formation of force-resistant kinetochore-microtubule attachments, explaining why deletion of the loop ablates chromosome congression. The loop may fold into a more rigid structure than previously assumed and we identify point mutations that impair Ndc80-Ndc80 interactions and recapitulate the effects of loop depletion. The loop also promotes spindle checkpoint signaling, suggesting that the organisation of adjacent Ndc80 complexes is crucial to couple kinetochore-microtubule binding to cell cycle control.

## INTRODUCTION

Establishing firm attachments between mitotic chromosomes and spindle microtubules is essential for faithful chromosome segregation and genome stability. Kinetochores, large protein platforms that assemble on centromeric DNA, mediate the attachment of chromosomes to microtubules. During mitosis, kinetochores first bind the lateral surface of microtubules to support the congression of chromosomes towards the middle of the dividing cell. They then capture the ends of dynamic microtubules to promote chromosome biorientation on the mitotic spindle (Tanaka et al., 2005; Kapoor et al., 2006; Akiyoshi et al., 2010; Magidson et al., 2011; Shrestha and Draviam, 2013; Chakraborty et al., 2019).

The widely conserved Ndc80 heterotetramer is the prime mediator of the binding between kinetochores and the ends of microtubules (Cheeseman et al., 2006; DeLuca et al., 2006; Ciferri et al., 2008; Wei et al., 2007; Guimaraes et al., 2008; Alushin et al., 2010; Sundin et al., 2011; Tooley et al., 2011). Ndc80 binds microtubules through the N-terminal calponin homology (CH) domains of its NDC80 (also named HEC1) and NUF2 subunits, with a contribution of the unstructured NDC80 N-terminal tail. It docks onto the inner kinetochore through the RWD domains of its SPC24 and SPC25 subunits. The CH- and RWD-domains are separated by a long coiled-coil stalk (Musacchio and Desai, 2017).

The spindle assembly checkpoint (SAC) delays anaphase onset until all chromosomes have bioriented on the mitotic spindle. Various mitotic kinases and phosphatases control the interplay between chromosome biorientation and SAC signaling. A crucial aspect of this dynamic interplay is the ability to identify and correct erroneous (e.g. syntelic) kinetochore-microtubule attachments. Key signaling substrates are well known (for example KNL1 and the NDC80 tail), but how they respond to kinetochore-microtubule binding in general, and to biorientation specifically, is poorly understood (Saurin, 2018).

Tension exerted by spindle microtubules pulling on kinetochores has a crucial role in error correction (Lampson and Grishchuk, 2017), but molecular cues for tension signaling at kinetochores remain elusive. As an elongated microtubule-binder in the outer kinetochore, the Ndc80 complex is a prime candidate to sense and signal tension. One hypothesis is that microtubule-bound Ndc80 complexes undergo a tension-dependent conformational change from a bent to a stretched state (the “jackknife” model) (Scarborough et al., 2019; Roscioli et al., 2020). In turn, tension-dependent conformational changes in the Ndc80 complex may control docking sites for kinases, phosphatases, or additional microtubule binders. Ndc80 bending in the jackknife model has been proposed to require the Ndc80 loop, a previously identified sequence in the NDC80 subunit predicted to interrupt the continuous coiled coil (Wigge et al., 1998). Studies in budding yeast, fission yeast, and humans demonstrated that the Ndc80 loop is essential for the establishment of end-on kinetochore-microtubule attachment and proper chromosome segregation (Hsu and Toda, 2011; Maure et al., 2011; Zhang et al., 2012; Tang et al., 2013; Shrestha and Draviam, 2013; Wimbish et al., 2020).

In human cells, a bioriented kinetochore contains approximately 250 copies of the Ndc80 complex (Suzuki et al., 2015) and binds a bundle of approximately 10 microtubules, known as a k-fiber (O’Toole et al., 2020; Kiewisz et al., 2022). This predicts that a single k-fiber microtubule may be bound by multiple Ndc80 complexes (**Figure 1A**). Reinforcing this prediction, single Ndc80 complexes fail to bind to the ends of dynamic microtubules *in vitro*, but artificial multimerisation overcomes this and generates robust end-tracking (Powers et al., 2009; Volkov et al., 2018), thus suggesting a crucial role of Ndc80 multivalency. From a mechanistic perspective, however, it remains unclear whether the acquisition of end-tracking reflects the uncoordinated binding of multiple individual Ndc80 complexes to the microtubule, or instead reflects additional cooperative interactions between Ndc80 complexes elicited by microtubule binding. The existing evidence is conflicting. In the absence of microtubules, Ndc80 complexes do not appear to be able to form higher oligomeric structures (Huis in ‘t Veld et al., 2016), but clustering of Ndc80 complexes has been observed along a microtubule lattice (Ciferri et al., 2008; Alushin et al., 2010, 2012). Neighboring Ndc80 complexes have been proposed to align upon microtubule binding (Yoo et al., 2018; Roscioli et al., 2020), thus suggesting that they interact laterally. Conversely, a modeling study suggested that a “lawn” of non-interacting Ndc80 complexes is ideally suited to generate robust microtubule binding (Zaytsev et al., 2015).

**Figure 1.**
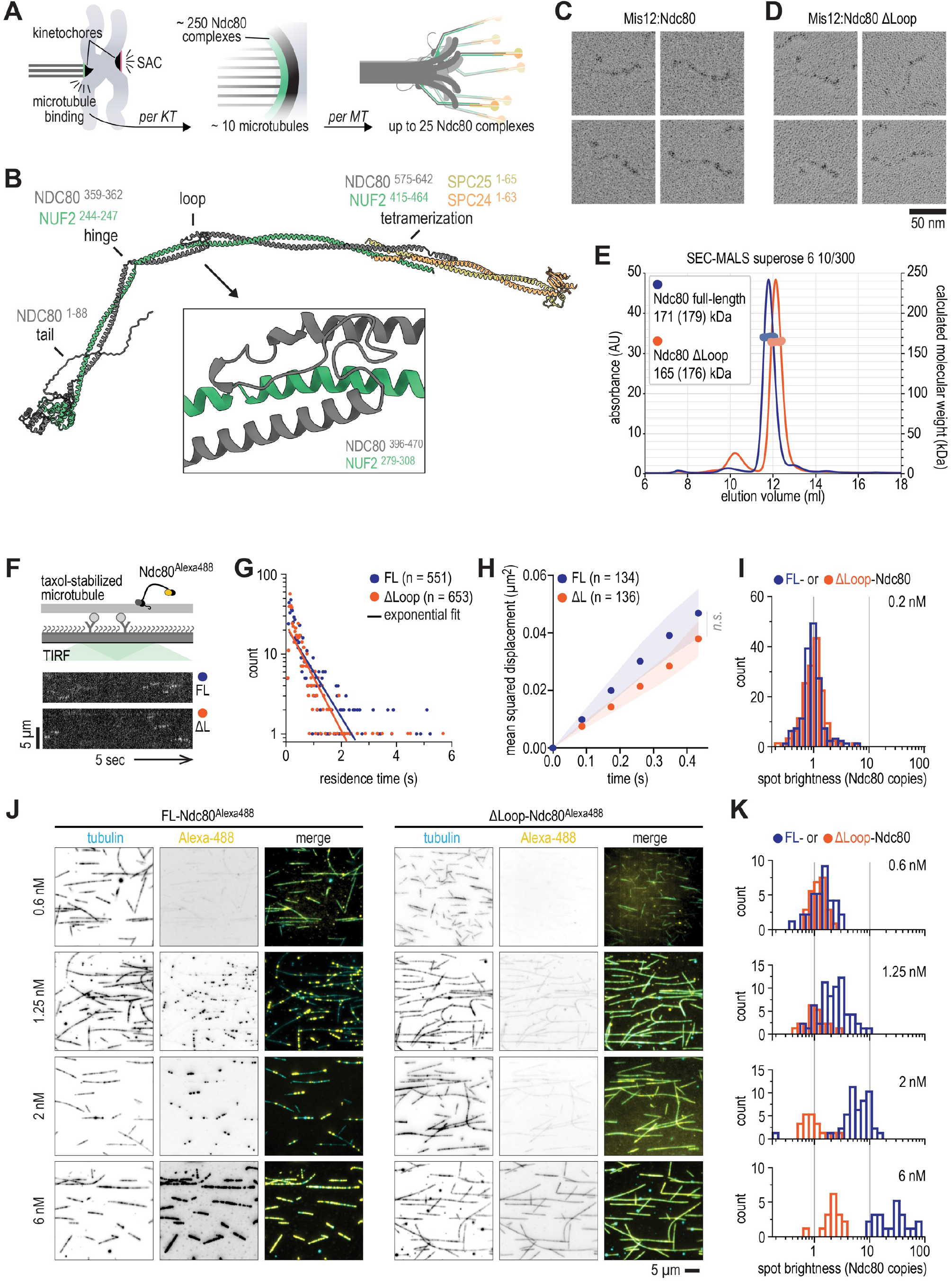
Structural analysis and loop-dependent clustering of Ndc80 on microtubules. **A**) Cartoon of a chromosome pair during mitosis with sister kinetochores that are attached (green) and unattached (red) to microtubules. The unattached kinetochores trigger a spindle assembly checkpoint (SAC) signal. One outer kinetochore contains a lawn of Ndc80 complexes, resulting in many Ndc80 complexes binding to a single microtubule. **B**) Prediction of the full-length Ndc80 structure with residues that comprise the tail, th e kink, the loop, and the tetramerisation domain indicated. The box shows the loop region at a 6x magnification. See **Suppl. Figure 1** for more information. **C, D**) Low-angle Pt/C shadowing of Mis12:Ndc80 (panel C) and Mis12:Ndc80^Δloop (Δ431-463)^ (panel D) complexes. The Mis12 complex appears as a 20 nm rod-like extension and marks Ndc80 and Ndc80^Δloop^. Calculated and (theoretical) masses are indicated in the legend. These complexes were fluorescently labelled. See **Suppl**. the SPC24:SPC25 side of the Ndc80 complex. **E**) Size exclusion chromatography coupled with multi-angle light scattering (SEC-MALS) profiles of **Figure 3** for more information. **F**) Total Internal Reflection Fluorescence (TIRF) microscopy was used to investigate Ndc80^Alexa488^ complexes on taxol-stabilized microtubules that were attached to a passivated glass surface. Kymographs show Ndc80 complexes at a concentration of 0.2 nM with (FL, blue) or without (ΔL, orange) the loop. Scale bars: vertical (5 μm), horizontal (5 s). **G**) Quantification of Ndc80 residence times for data as in panel F. Solid line represents a single exponential fit. **H**) One-dimensional diffusion of Ndc80 complexes (with *n* indicated) on microtubules. Traces were split into segments of 0.5 s and averaged (see Materials and Methods). Mean values (circles) and SEM (shaded area) are shown. **I**) Distribution of the initial brightness of Ndc80 complexes on stabilized microtubules. **J**) Typical fields of view showing decoration of taxol-stabilized microtubules (cyan) incubated with full-length or loopless Ndc80 (yellow) at the indicated concentration. Images show an average projection of 200 frames. The contrast between individual fluorescent channels (inverted grayscale) was fixed. Auto-contrast was used for the composite images to highlight the differences in the uniformity of Ndc80 decoration. **K**) Distribution of the initial brightness of Ndc80 complexes added at indicated concentrations to taxol-stabilized microtubules.

Here we demonstrate that direct interactions between full-length Ndc80 complexes on microtubules are essential for the formation of stable kinetochore-microtubule attachments in a fully reconstituted system and in dividing cells. These interactions are perturbed by deletion of the Ndc80 loop, and by point mutations therein. Our study provides structural insight into the Ndc80 loop, a mechanistic explanation for futile chromosome congression cycles in loop mutants, and suggests a direct contribution of the Ndc80 loop to SAC signaling.

## RESULTS

### The Ndc80 loop folds rigidly against NUF2 and NDC80

To gain structural understanding of Ndc80, we predicted the structure of full-length Ndc80 using AlphaFold-Multimer (Evans et al., 2022) (**Figure 1B**). Except for the unstructured tail of the NDC80 subunit, the structure of full-length Ndc80 showed high to very high confidence scores (**Suppl. Figure 1**), thus accurately predicting the relative positions of NDC80, NUF2, SPC25, and SPC24, including in the tetramerisation region in which the coiled coils of all four subunits overlap (**Suppl. Figure 1**) (Valverde et al., 2016). Very high confidence scores were also attributed to the Ndc80 loop, a region comprising residues 421-450 of the NDC80 subunit that interrupts the long NDC80:NUF2 coiled coil. In the predicted structure, the loop connects two slightly staggered alpha-helices of NDC80, both of which pair with the continuous NUF2 alpha-helix (**Figure 1B**). The loop makes multiple turns, but does not bulge out freely from the helical axis of the complex as previously speculated (*e*.*g*. (Ciferri et al., 2008)). Rather, it uses conserved amphipathic stretches to pack closely and rigidly against the NUF2 and NDC80 helices. We hypothesize that two strongly conserved cysteine residues, NUF2^C289^ and NDC80^C449^, might further stabilize this structure through a disulfide bond. The loop region itself spans 30 residues in humans and other higher eukaryotes, but exceeds 60 residues in other species, indicating remarkable variation of composition and length.

The loop of some species, such as *Candida albicans* and *Dictyostelium discoideum*, contain low-complexity regions with stretches of hydrophilic residues. Despite these remarkably divergent sequences, the core folds of the predicted loop regions were comparable (**Suppl. Figure 2**). Thus, the loop is unlikely to be a point of bending along the Ndc80 coiled-coils. Rather, a prominent hinge in the Ndc80 structure is predicted upstream from the loop, around residues NDC80^359-362^ and NUF2^244-247^, (**Figure 1B**). The hinge is conserved across eukaryotes and its position is in agreement with micrographs of full-length Ndc80 complexes (Wang et al., 2008; Huis in ‘t Veld et al., 2016; Jenni and Harrison, 2018) (**Figure 1C** and **Suppl. Figure 2**).

**Figure 2.**
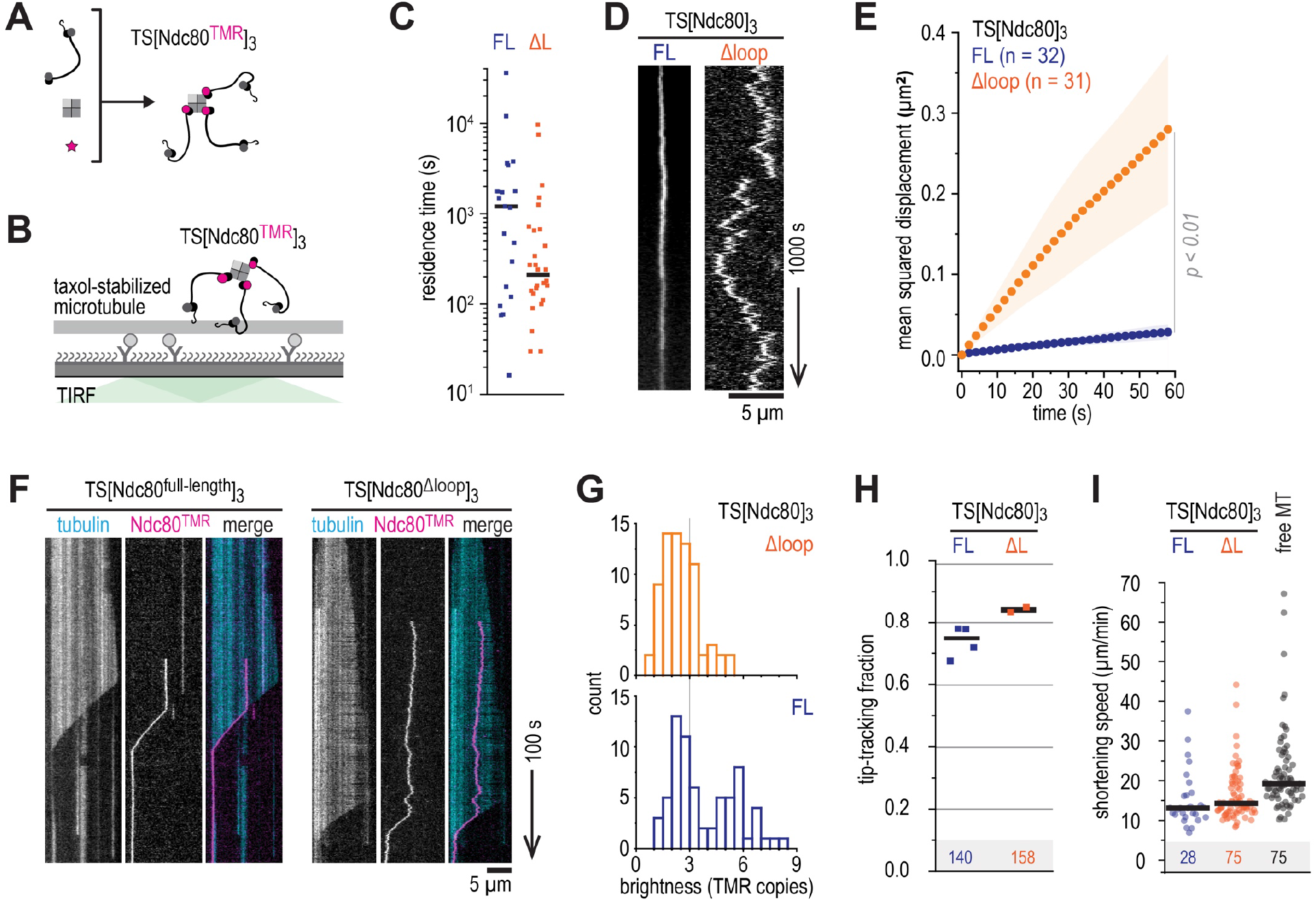
The loop reduces diffusion of Ndc80 trimers without affecting their end-tracking. **A**) Preparation of TMR-labelled, streptavidin-mediated Ndc80 trimers. See **Suppl. Figure 3** for more information. **B, C, D**) TMR-labelled Ndc80 trimers with (FL, blue) or without (ΔL, orange) the loop were added to taxol-stabilized microtubules to measure their residence time (panel C) and one-dimensional diffusion (panel D). Ndc80 trimers with and without the loop are shown. Scale bars: vertical (1000 s), horizontal (5 μm). **E**) One-dimensional diffusion of Ndc80 trimers (with *n* indicated) on microtubules. Traces were split into segments of 60 s and averaged (see Materials and Methods). Mean values (circles) and SEM (shaded area) are shown. **F**) Kymographs of full-length and loopless Ndc80 trimers (10 pM) that reside on dynamically growing and shortening microtubules. Trimers remain bound to the ends of shortening microtubules. Scale bars: vertical (100 s), horizontal (5 μm). **G**) Distribution of initial brightness of end-tracking Ndc80 trimers with full-length (blue) or loopless (orange) Ndc80. **H**) Fraction of Ndc80 trimers that switches from lateral microtubule binding to tracking shortening ends. Data from four repeats (total 140 events) for full-length Ndc80 trimers (blue), and two repeats (total 158 events) for loopless Ndc80 trimers (orange). Horizontal lines indicate average values. **I**) End-tracking speed of full-length (blue) and loopless (orange) Ndc80 trimers that follow shortening microtubule ends. Compared to Ndc80-free shortening ends in the same field of views (gray). Horizontal lines indicate median values.

To investigate the functional and structural relevance of the loop, we generated a recombinant human Ndc80 complex lacking NDC80 residues 431-463. Full-length and loopless Ndc80 complexes (in complex with the Mis12 complex to distinguish its ends) were essentially indistinguishable (**Figure 1C-D**). While the hinge was a prominent point of bending, both complexes displayed additional bending points along their length. SEC-MALS and mass photometry further supported that full-length and loopless Ndc80 complexes have comparable structural and biochemical properties (**Figure 1D** and **Suppl. Figure 3A-C**). Using DSBU, a crosslinker that preferentially targets primary amines, but also reacts with proximal hydroxyls, we identified 190 unique crosslinks by mass spectrometry (**Suppl. Figure 4**). Most crosslinks connect side-chains that are less than 30 Å apart in our predicted structure and are consistent with previous analyses (Maiolica et al., 2007; Pan et al., 2018; Helgeson et al., 2018; Huis in ‘t Veld et al., 2019). High-scoring crosslinks are predominantly found near the tetramerisation domain and the Ndc80 loop and provide experimental support for the structural prediction (**Suppl. Figure 4**).

**Figure 3.**
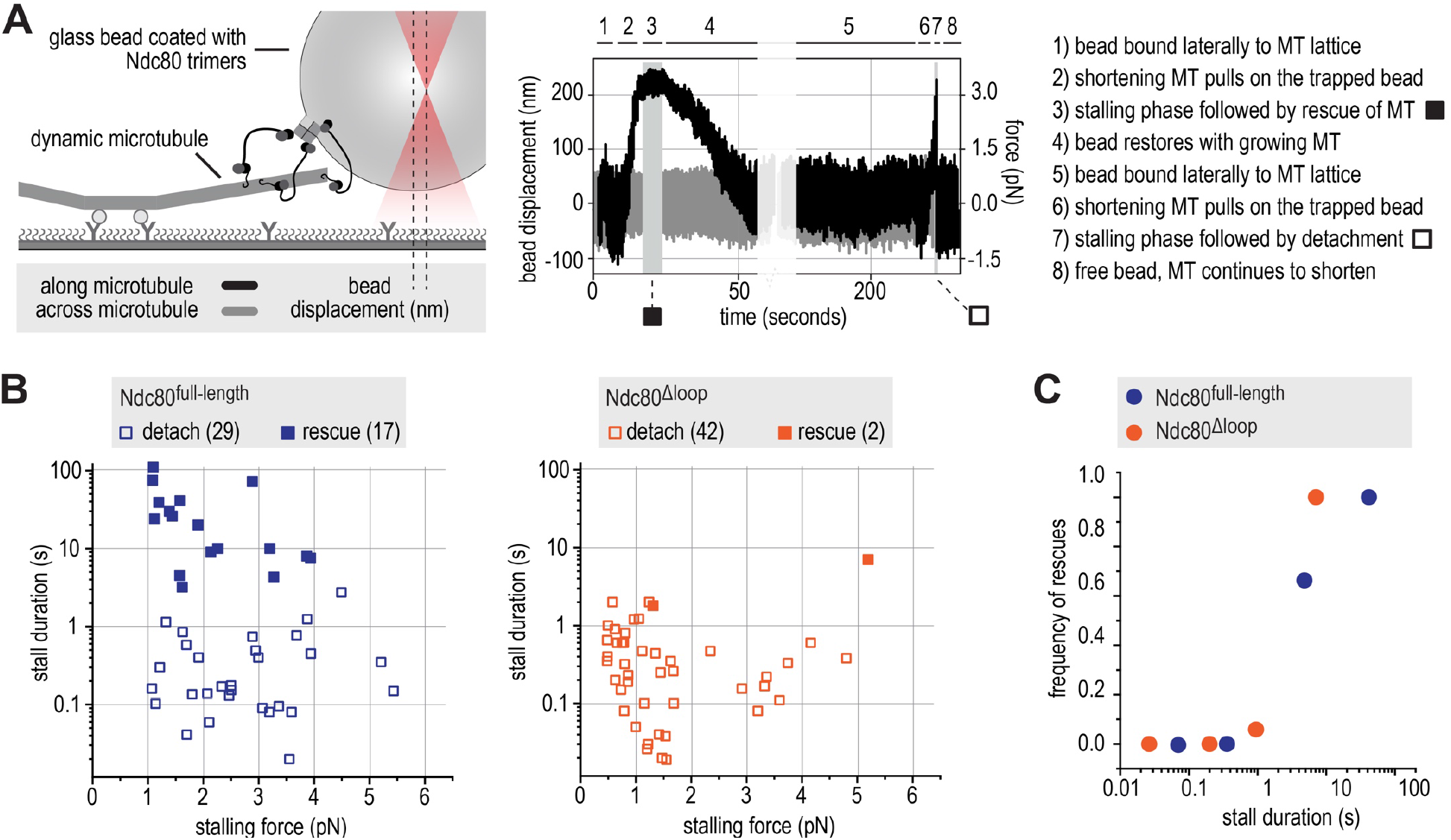
The loop stabilizes end-on Ndc80-microtubule interactions under force. **A**) Schematic of the optical trap experiment and a typical force trace. A glass bead coated with full-length or loopless Ndc80 trimers is held in an optical trap near a microtubule end. The displacement of the bead (left Y axis) and the corresponding force (right Y axis) are shown along and across the microtubules axis (black and grey, respectively). Typical stages of an experiment are described in the legend. **B**) Correlations of stall duration and stall force. Each datapoint represents a single stall event. Filled squares: stalls resulting in microtubule rescue. Open squares: stalls resulting in bead detachment from the microtubule. **C**) The frequency of rescue events after a stall are plotted for full-length (*n* = 46) and loopless (*n* = 44) Ndc80 after binning based on stall duration.

**Figure 4.**
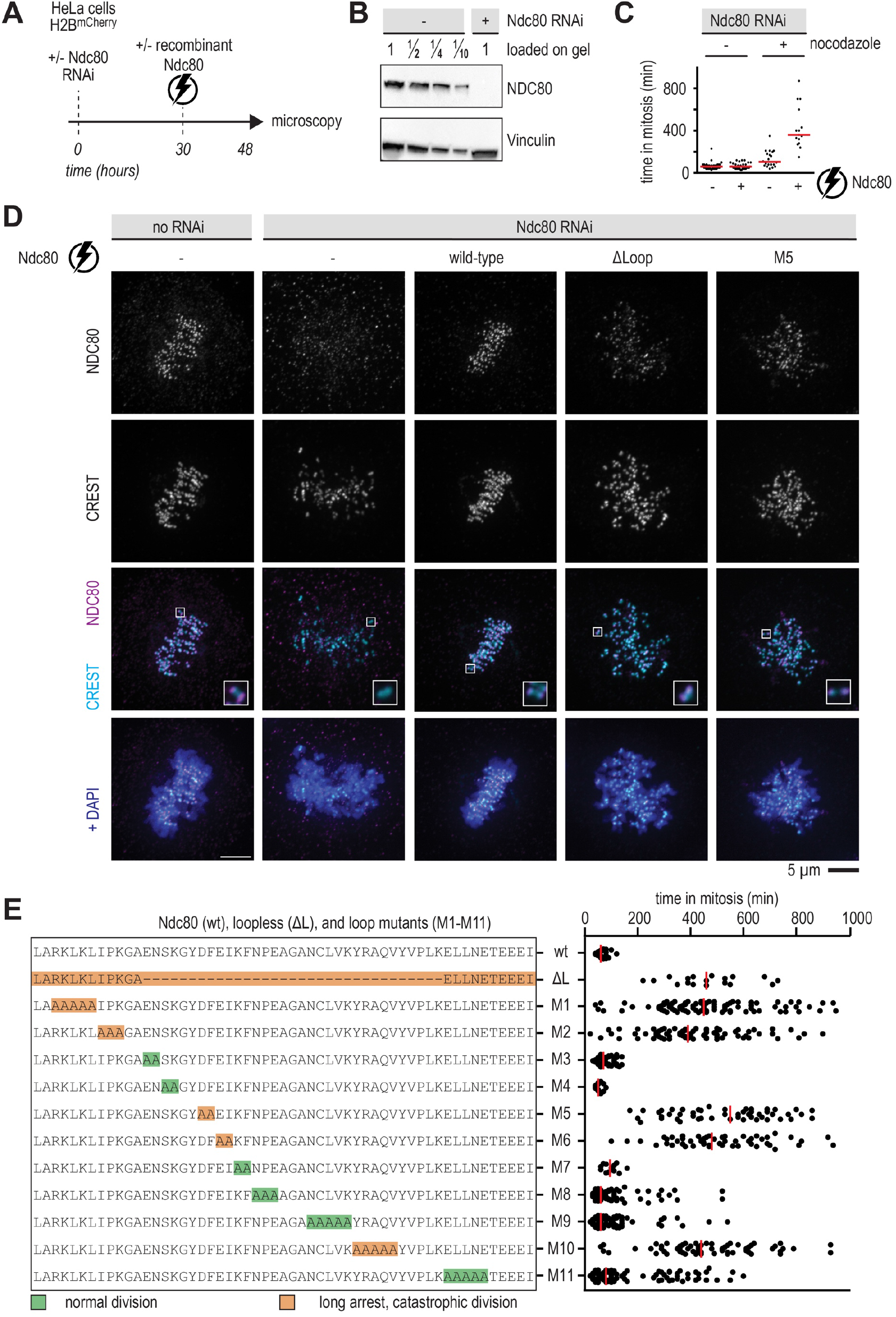
Mutation of critical residues in the loop impairs chromosome congression. **A**) Schematic of an electroporation experiment. **B**) Immunoblot of NDC80 levels following depletion of the Ndc80 complex by RNAi. **C**) Quantification of the time that cells spent in mitosis following various treatments. Each dot represents a cell and the red lines indicate median values. Nocodazole was added 17 hours after electroporation and 1 hour before microscopy. A minimum of 30 cells were analysed for each condition. **D**) Immunofluorescence microscopy of mitotic cells stained for DNA (DAPI), kinetochores (CREST), and Ndc80 complexes. The Ndc80 antibody (9G3) detects endogenous Ndc80 (column 1) and electroporated recombinant Ndc80 (columns 3, 4, 5). Representative cells with a metaphase plate or uncongressed chromosomes are shown. Scale bar: 5 μm. **E**) Overview of mutations in the Ndc80 loop region and the effects on the time spent in mitosis following the experimental setup outlined in panel A. Colors indicate whether cells divided normally, sometimes with delayed chromosome congression (green), or showed long arrests followed by a catastrophic division (orange). Each dot represents a cell and the red lines indicate median values. Mutation NDC80^G434A-Y435A^ did not support the formation of stable and soluble Ndc80 complexes. A minimum of 30 cells were analysed for each condition.

### Ndc80 complexes cluster on the microtubule lattice in a loop-dependent manner

To test if the loop directly affects the binding of Ndc80 to microtubules, we labelled recombinant full-length and loopless Ndc80 with AlexaFluor^488^ using Sortase (Hirakawa et al., 2015) and studied their binding to stabilized microtubules (**Figure 1F** and **Suppl. Figure 3A-C**). At sub-nanomolar concentrations (0.2 nM), full-length and loopless Ndc80 complexes typically resided on microtubules for 0.2 - 2 seconds with one-dimensional diffusion (D) along the microtubule lattice of 0.10 and 0.09 μm^2^/s, respectively (**Figure 1G-I**). These values are in agreement with previously reported values for truncated Ndc80 bonsai complexes (Zaytsev et al., 2015) and budding yeast Ndc80 (Powers et al., 2009; Scarborough et al., 2019).

Full-length and loopless Ndc80 complexes, however, bound microtubules in distinct ways at higher concentrations. Specifically, full-length Ndc80 formed clusters on the microtubule lattice whereas loopless Ndc80 did not. The clusters increased from 3-6 Ndc80 complexes at 1.25 nM, to 5-10 complexes at 2 nM, and to 10-100 complexes at 6 nM (**Figure 1J-K**). Full-length Ndc80 thus appears to bind microtubules preferably near sites already occupied by Ndc80, presumably through interactions between adjacent Ndc80 complexes. Since loopless Ndc80 binds uniformly along microtubules, we conclude that clustering of Ndc80 complexes upon microtubule binding requires an intact Ndc80 loop (**Figure 1J-K**).

### Loop-dependent coordination of Ndc80-microtubule binding

We previously used modules with three Ndc80 arms coupled to an engineered streptavidin-derived scaffold to mimic the modular organisation of Ndc80 in the outer kinetochore (Volkov et al., 2018; Huis in ‘t Veld et al., 2019). To understand the contribution of loop-mediated Ndc80-Ndc80 interactions in this context, we generated trimers of full-length and loopless Ndc80 and studied their interactions with microtubules *in vitro* (**Figure 2A-B** and **Suppl. Figure 3D**). Full-length and loopless Ndc80 trimers resided stably on microtubules with residence times ranging from minutes to hours (medians of 1000 and 200 seconds, respectively) (**Figure 2C**). Compared to the high diffusion rates of individual Ndc80 complexes on microtubules (**Figure 1H**), diffusion of a scaffold with three full-length Ndc80 complexes was greatly reduced (D = 4.3 ± 1.0 × 10^−4^ μm^2^/s), likely reflecting the presence of multiple microtubule-binding elements in Ndc80 trimers. Interestingly, the Ndc80 loop contributed significantly to this reduction, because trimers of loopless Ndc80 diffused an order of magnitude faster (D = 3.6 ± 1.6 × 10^−3^ μm^2^/s) (**Figure 2D-E**). Despite this difference in diffusion rates, loopless Ndc80 trimers efficiently tracked the ends of shortening microtubules and reduced microtubule depolymerisation rates, as previously observed for full-length Ndc80 trimers **(Figure 2F-I**) (Volkov et al., 2018; Huis in ‘t Veld et al., 2019). Thus, deletion of the Ndc80 loop reduces the stability of the interaction between Ndc80 trimers and the microtubule lattice, but does not grossly affect recognition of the microtubule plus end in the absence of applied forces.

### The loop is needed to stabilize end-on Ndc80-microtubule interactions under force

Next, we set out to compare how full-length and loopless Ndc80 bind to the ends of shortening microtubules under force. We coated biotinylated glass beads with Ndc80 trimers and moved them with an optical tweezer to the lattice of a dynamic microtubule. When the microtubule started to depolymerize, we measured the forces exerted by the shortening end of the microtubule against the trapped bead (**Figure 3A**). The microtubule-generated force typically increased until it matched the opposing force generated by the displacement of the bead from its rest position in the optical trap, a condition that stalled microtubule shortening. A stall was either followed by a detachment, or by a switch of the microtubule to a growing state (rescue). Detached beads rapidly moved back to the centre of the trap. When a rescue allowed the bound microtubule to resume growth, beads retaining attachment to the microtubule moved back to the trap centre much more gradually (**Figure 3A**).

Full-length Ndc80 trimers often stalled microtubule depolymerisation for seconds and rescued microtubule growth in 17 out of 46 cases (37%) (**Figure 3B**), in agreement with our previous experiments (Volkov et al., 2018; Huis in ‘t Veld et al., 2019). By contrast, beads coated with loopless Ndc80 trimers only rescued microtubule growth in 2 out of 44 cases (5%) and typically detached from shortening microtubules after stalls that were usually shorter than one second (**Figure 3B**). We showed previously that stalls of microtubule depolymerisation that last more than approximately one second are much more likely than shorter stalls to result in a rescue of microtubule growth, even with stall forces as low as 1 pN (Huis in ‘t Veld et al., 2019). This relation was also clearly observed with our new measurements, as the majority of stalls that led to rescues in presence of full-length Ndc80 lasted more than one second, whereas most stalls with loopless Ndc80 were short and followed by detachment (**Figure 3B-C**). Full-length Ndc80 trimers bound more efficiently than loopless trimers to the biotinylated glass beads, possibly because the loop-mediated Ndc80-Ndc80 interactions described above facilitate coating. Nonetheless, beads coated with high amounts of loopless Ndc80 also failed to efficiently stall and rescue shortening microtubules (**Suppl. Figure 3E-F**). Taken together, our single-molecule studies suggest that loop-mediated interactions coordinate Ndc80 complexes into a non-diffusive microtubule-binder that forms load-bearing attachments with the ends of microtubules. Since a single kinetochore contains many copies of Ndc80, the clustering of adjacent Ndc80 complexes might be important for the attachment of kinetochores to the dynamic microtubules of the mitotic spindle.

### The Ndc80 loop is essential for proper chromosome congression

To investigate how the loop influences kinetochore-microtubule interactions in mitotic cells, we electroporated recombinant Ndc80 complexes into HeLa cells stably expressing mCherry-H2B and depleted of endogenous Ndc80 by RNAi. This approach directly assesses if recombinant complexes with validated stoichiometry and stability support cell division (Alex et al., 2019). We used live-cell imaging to follow fluorescently labelled recombinant Ndc80 complexes 18 hours after they were delivered into cells (**Figure 4A**). Endogenous Ndc80 complexes were efficiently depleted upon exposure to a siRNA combination targeting NDC80, SPC25, and SPC24 (**Figure 4B**). Depletion of Ndc80 ablated the spindle assembly checkpoint (SAC) and caused cells to exit mitosis in presence of the microtubule poison nocodazole, in line with previous work that established a requirement of Ndc80 in SAC signaling (**Figure 4C**) (McCleland et al., 2003; Kim and Yu, 2015). When recombinant full-length Ndc80 complexes were delivered into cells depleted of endogenous Ndc80, they restored the expected cell cycle arrest in nocodazole-treated cells (**Figure 4C**). Collectively, these control experiments demonstrate the efficient depletion of endogenous Ndc80 and the functionality of the electroporated recombinant Ndc80 complex.

Using this assay, we established that both full-length and loopless Ndc80 complexes were efficiently recruited to kinetochores in cells that lack endogenous Ndc80. However, whereas full-length recombinant complexes supported the timely formation of a metaphase plate and faithful cell division (**Figure 4D**, 3^rd^ column), cells electroporated with loopless complexes failed to congress their chromosomes to the midplane and remained in a prometaphase-like state without ever fully forming a recognizable metaphase plate (**Figure 4D**, 4^th^ column). Thus, the loop region is essential for chromosome alignment and bi-orientation, as previously shown (Zhang et al., 2012; Shrestha and Draviam, 2013; Wimbish et al., 2020). In agreement with previous studies (Zhang et al., 2012; Wimbish et al., 2020), cells with loopless Ndc80 complexes arrested in mitosis for many hours (**Figure 4E**). This indicates that the loopless Ndc80 mutant is impaired in bi-orientation, but restores a contribution to SAC signaling that is instead lost upon Ndc80 depletion (as clarified in the previous paragraph).

To dissect the properties of the NDC80 loop in more detail, we generated full-length Ndc80 complexes with residues of the loop mutated into Alanine. A complex carrying the NDC80^G434A-Y435A^ mutations was insoluble, but the other eleven complexes were stable and were purified to homogeneity. After adding a small fluorescent dye, complexes were electroporated into cells depleted of endogenous Ndc80 complex (**Figure 4E**). Wild-type Ndc80 complexes and mutants 3, 4, 7 supported chromosome congression and segregation similarly to wild-type Ndc80. Mutants 8, 9, 11 displayed almost normal division, but delayed chromosome congression was often observed. Mutants 1, 2, 5, 6, 10, on the other hand, phenocopied the prometaphase-like arrest observed with loopless Ndc80, indicating that these mutations profoundly impair loop function (**Figure 4E**). The loop mutant NDC80^D436A-F437A^ (M5) phenocopied the deletion of the entire loop in a chromosome alignment assay (**Figure 4D**, 5^th^ column). Since a role of residues NDC80^436-439^ in chromosome congression was also reported in a previous study (Wimbish et al., 2020) (**Suppl. Fig 5A**), we chose this mutant for further analyses. The composition of the outer kinetochore, as assayed by quantifying the staining for components KNL1 and NSL1, was essentially identical in cells with Ndc80 full-length, loopless, or M5 (**Suppl. Figure 5B**). Moreover, kinetochores with different Ndc80 mutants recruited comparable amounts of SAC components in the presence of nocodazole (**Suppl. Figure 5B**).

**Figure 5.**
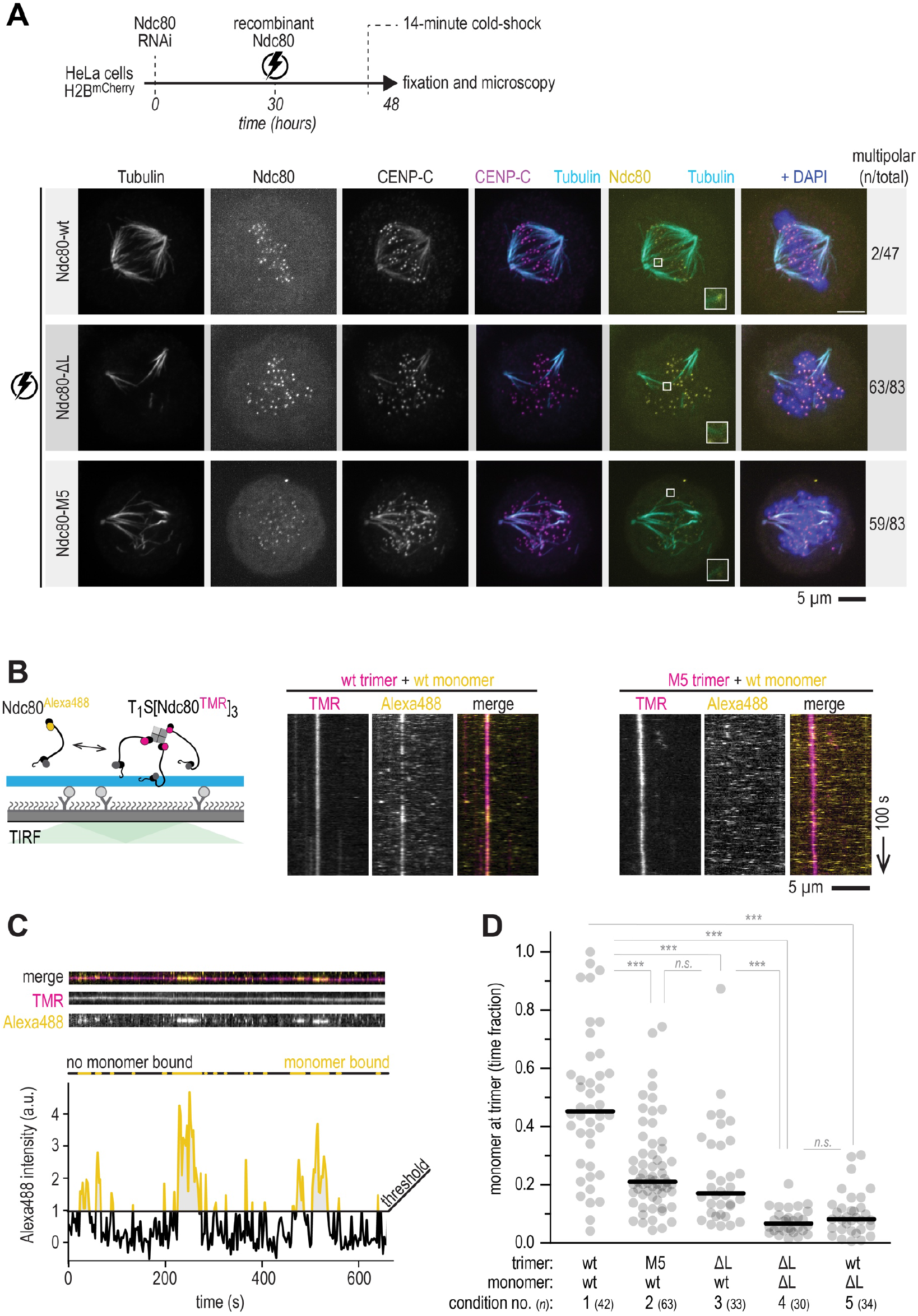
The M5 mutant phenocopies loopless Ndc80 *in vivo* and *in vitro*. **A**) Schematic of a cold-shock assay following an electroporation experiment. Immunofluorescence images showing the attachment status of kinetochores to microtubules in cells electroporated with recombinant Ndc80-wt, Ndc80-ΔL, or Ndc80-M5 complexes. The number of cells with multipolar spindles and the total number of analysed cells are shown. Some signal from the tubulin channel is visible in the CENP-C channel. Scale bar: 5 μm. **B**) Total Internal Reflection Fluorescence (TIRF) microscopy was used to investigate Ndc80^Alexa488^ complexes (0.6 nM) added to trimeric Ndc80^TMR^ (10 pM) on fluorescent taxol-stabilized microtubules that were attached to a passivated glass surface. Typical kymographs showing virtually motionless Ndc80 trimers (magenta) and transiently binding Ndc80 monomers (yellow). Wild-type (wt) monomers associate with wt trimers (left), but not with M5 trimers (right). Scale bars: vertical (100 s), horizontal (5 μm). **C**) Quantification of the intensity of the monomeric Ndc80 associating with microtubule-bound trimeric Ndc80 (see Materials and Methods for details). A threshold for binding was set at an intensity equivalent to one Alexa488 copy. Intensities well above 1 (yellow) could thus reflects multiple monomers binding simultaneously. **D**) Fraction of time there was at least one monomer (added to solution at a concentration of 0.6 nM) present at the microtubule-bound trimer (10 pM), tested in various combinations of wild-type (wt), loopless (ΔL) and M5 monomers or trimers. All analysed traces of Ndc80 trimers are shown (*n* is shown in the legend). Horizontal lines show median values and statistical significance was determined using a two-tailed Mann-Whitney test. P-values: 1 (wt-trimer + wt-monomer) vs 2 (M5-trimer + wt-monomer): 7·10^−7^ (***); 2 vs 3: 0.17 (n.s.); 1 vs 3: 1·10^−6^ (***); 1 vs 4: 3·10^−15^ (***); 3 vs 4: 1·10^−8^ (***); 1 vs 5: 1·10^−13^ (***); 4 vs 5 0.23 (n.s.).

### The M5 mutant phenocopies loopless Ndc80

To further test how the Ndc80 loop promotes kinetochore-microtubule interactions *in vivo*, we exposed cells electroporated with wild-type or mutated Ndc80 to a cold shock before fixation for immunofluorescence microscopy (**Figure 5A**). Properly formed metaphase plates and cold-stable kinetochore-microtubule fibers were observed in cells with wild-type Ndc80, but not in cells with loop mutants (**Figure 5A**). The frequent observation of multipolar spindles in cells with Ndc80 loop mutations supports the idea that the loop is required for stable kinetochore-microtubule interactions. To further investigate the propensity of Ndc80 to form clusters, we measured the association of monomeric full-length or loopless Ndc80^Alexa488^ with Ndc80^TMR^ trimers already bound to microtubules (**Figure 5B**). Wild-type Ndc80 monomers (at a concentration of 0.6 nM) interacted with microtubules markedly longer when they colocalized with wild-type Ndc80 trimers, suggesting an interaction that prolongs their residence on microtubules. Monomers of loopless Ndc80, on the other hand, were indifferent to the presence of wild-type trimers, and resided on microtubules for the same short time observed when wild-type monomers bound microtubules in the absence of trimers (**Suppl. Fig 6**). We then analysed the microtubule residency time of various combinations of wild-type or mutant trimeric Ndc80^TMR^ and monomeric Ndc80^Alexa488^ (**Figure 5B-C**). Wild-type trimers bound wild-type monomers for 48 ± 25% (mean ± SD) of the observation time, significantly longer than M5 trimers (25 ± 15%) and loopless trimers (23 ± 17%) (**Figure 5D** and **Suppl. Fig 6**). This fraction was further reduced for loopless monomers recruited to wild-type trimers (10 ± 8%), or to loopless trimers (7 ± 4%). These experiments indicate that residues NDC80^D436-F437^, which are mutated in the M5 mutant, directly contribute to Ndc80-Ndc80 interactions on microtubules.

**Figure 6.**
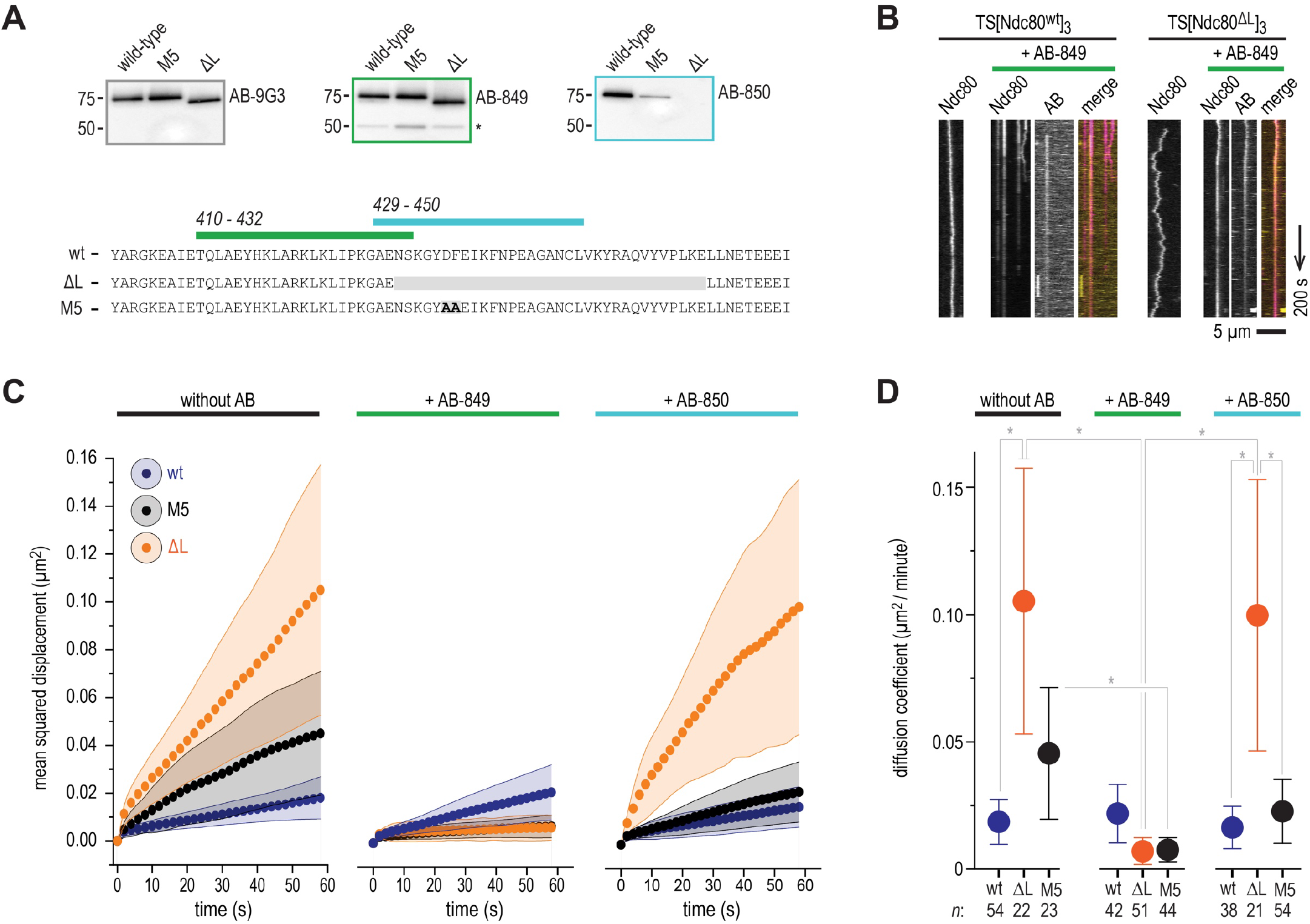
Loop-proximal Ndc80-Ndc80 crosslinking rescues increased diffusivity of loopless Ndc80 trimers. **A**) Representation of the peptides used to raise AB-849 and AB-850 and immunoblots showing their recognition of wild-type, M5 and loopless Ndc80 complexes. 9G3 is a commercially available monoclonal antibody raised against NDC80^56-642^, later shown to recognize NDC80^200-215^. Asterisk shows the non-specific recognition of another protein, presumably NUF2. **B**) Diffusion of full-length and loopless Ndc80 trimers in absence and presence of AB-849. The primary rabbit polyclonal AB-849 was detected using a Alexa650-labelled anti-rabbit secondary IgG antibody. Scale bars: vertical (100 s), horizontal (5 μm). **C**) One-dimensional diffusion of full-length (blue), loopless (orange), and M5 (black) Ndc80 trimers in presence and absence of AB-849 and AB-850. Traces of Ndc80 trimers (with *n* indicated in the legend for panel D) on microtubules were analysed. Traces were split into segments of 60 s and averaged (see Materials and Methods). Mean (circles) and SEM (shaded areas) values are shown. We note that the omission of reducing agents, a precondition to use the antibody as a crosslinker, slightly decreased the overall diffusion of Ndc80-modules on microtubules (compare with **Figure 2E**). **D**) Summary showing diffusion coefficients (μm^2^/minute) that follow from the data shown in panel C. Mean values, SEM, and number of diffusion traces (*n*) are indicated. Statistically significant differences were determined using a two-tailed t-test. P-values: no AB, FL vs ΔL: 0.0180 (*); ΔL, no AB vs AB-849: 0.0108 (*); M5, no AB vs AB-849: 0.0458 (*); ΔL, AB849 vs AB-850: 0.0156 (*); AB-850, ΔL vs FL: 0.0461 (*); AB-850, ΔL vs M5: 0.0494 (*).

### Loop-proximal Ndc80-Ndc80 crosslinking can rescue loop deletion *in vitro*

Our experiments demonstrated that multivalent Ndc80 requires the loop to form load-bearing attachments to microtubule-ends *in vitro* and *in vivo*, presumably by clustering individual microtubule-binding elements into a robust microtubule-binding unit. To test this model further, we set out to crosslink Ndc80 arms artificially using polyclonal antibodies that were raised against short regions positioned just before or directly in the loop (NDC80^410-432^/AB-849 and NDC80^429-450^/AB-850, respectively) (**Figure 6A**). In the absence of Ndc80, neither of these antibodies interacted with microtubules *in vitro* (**Suppl. Fig 7A**).

**Figure 7.**
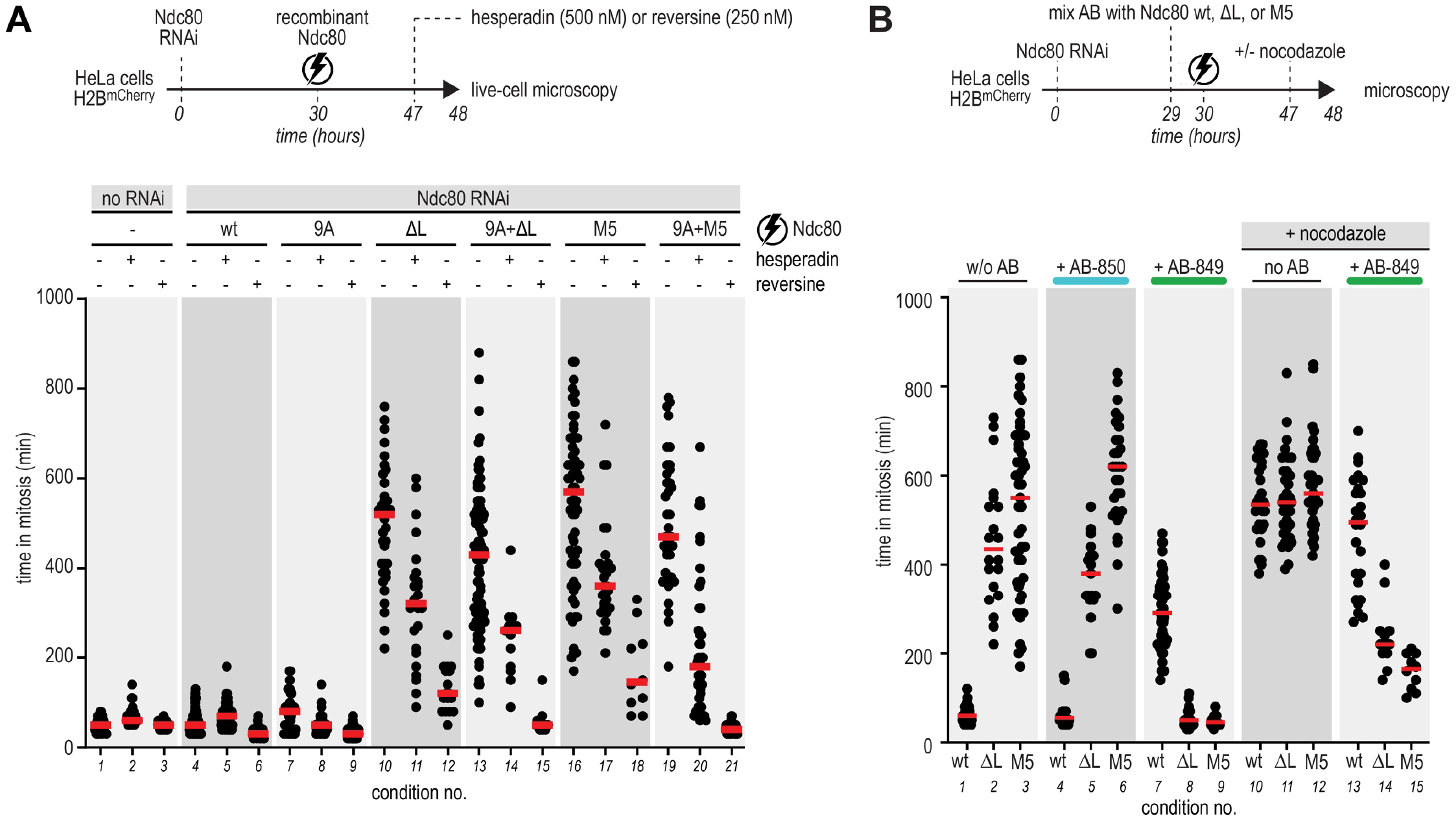
Loop-mediated Ndc80-Ndc80 interactions contribute to microtubule binding and SAC-signaling. **A**) Experimental workflow to investigate the time that cells, electroporated with recombinant Ndc80 complexes, spent in mitosis in the presence and absence of mitotic kinases inhibitors. Every dot represents a cell and red lines indicate median values. At least 30 cells were analysed for each condition. **B**) Experimental workflow to investigate how cells electroporated with various combinations of antibody (AB) and Ndc80 progress through mitosis. Every dot represents a cell and red lines indicate median values. At least 30 cells were analysed for each condition.

To test these antibodies and their putative effect on the clustering of Ndc80, we investigated their effect on the diffusion of Ndc80 trimers along microtubules. We first used a fluorescently labelled secondary antibody to confirm that the loop-directed antibodies bind Ndc80 modules on microtubules (**Figure 6B**) and experimentally determined the amount of primary antibody needed to bind Ndc80 trimers without causing antibody-induced aggregation. The fluorescence intensity of TMR-labelled Ndc80 was used to exclude multimers with more than one Ndc80 trimer from further analysis (**Suppl. Figure 7B-C**). The addition of AB-849, which binds close to the loop region, to loopless trimers greatly reduced their diffusion on microtubules. Such an effect was not observed with AB-850, which binds within the loop, and is therefore unable to bind loopless Ndc80 (**Figure 6C-D**). This experiment demonstrates that Ndc80-Ndc80 crosslinking near the loop is sufficient to reduce the diffusion of Ndc80 ensembles on microtubules.

### Loop-mediated mitotic arrest involves multiple phospho-signaling pathways

To prevent chromosome mis-segregation, a dividing cell needs to destabilize occasional syntelic and merotelic kinetochore-microtubule misattachments. This error correction process requires Aurora B, a kinase with multiple substrates in the kinetochore (Lampson and Grishchuk, 2017). Key Aurora B substrates include several serine and threonine residues in the 80-residue unstructured N-terminal tail of the NDC80 subunit, whose phosphorylation renders kinetochore-microtubule attachment reversible and is critical for error correction (Krenn and Musacchio, 2015). Inhibition of multi-site phosphorylation of the Ndc80 tail results in hyperstable kinetochore-microtubule attachments (DeLuca et al., 2006; Guimaraes et al., 2008).

We asked if preventing phosphorylation of the NDC80 tail had the potential to override the mitotic arrest caused by mutations in the loop region. We therefore combined nine Alanine (9A) mutations in the tail (preventing its phosphorylation) with either loopless or M5 mutants. Although the 9A Ndc80 tail reduced the duration of the mitotic arrest in loop mutants (**Figure 7A**, compare conditions 10 with 13 and 16 with 19), the double mutants sustained a robust mitotic arrest that lasted for several hours. Thus, artificially stabilizing microtubule attachment by individual Ndc80 complexes does not rescue the defect caused by tampering with the Ndc80 loop. This result suggests that Ndc80 clustering has a more fundamental role in establishing stable connections with microtubules, possibly one that precedes an assessment of the quality of the attachment by the error correction pathway.

In addition to controlling the error correction pathway, Aurora B also contributes to the spindle assembly checkpoint (SAC) response (Krenn and Musacchio, 2015). Predictably, addition of Hesperadin, a small molecule inhibitor of Aurora B kinase (Hauf et al., 2003), resulted in a severely shortened mitotic arrest in cells electroporated with Ndc80 loop mutants (**Figure 7A**, compare conditions 10 with 11 and 16 with 17). This shortening was much more significant than that caused by the Ndc80 9A error correction mutant, suggesting partial SAC abrogation, in addition to overriding the error correction pathway. This interpretation was corroborated by testing the effects of Reversine, a specific and potent inhibitor of the SAC kinase MPS1 (Santaguida et al., 2010). Reversine dramatically shortened the time that loop-deficient cells spent in mitosis (**Figure 7A**, conditions 12 and 18). This effect was exacerbated by the 9A mutation, likely because the enhanced microtubule binding by this mutant further facilitated checkpoint overriding by Reversine (**Figure 7A**, compare conditions 12 with 15 and 18 with 21). This result therefore suggests that the 9A mutant may not simply satisfy the SAC (Etemad et al., 2015; Tauchman et al., 2015), but also contribute to SAC overriding in the presence of microtubules.

### The Ndc80 loop contributes to SAC signaling

Finally, we decided to use AB-849 and AB-850 to examine the effects of antibody-mediated crosslinking on chromosome alignment and segregation. To do so, we incubated antibodies and recombinant Ndc80 complexes for one hour and electroporated the mixture into HeLa cells depleted of endogenous Ndc80 complex. Following pre-incubation of antibody and protein, AB-849 and AB-850 localized to kinetochores (**Suppl. Figure 8A**). AB-850, which binds the Ndc80 loop, did not appear to cause significant upheavals, as the pattern of mitotic arrest in cells previously depleted of endogenous Ndc80 complex and electroporated with wild-type, loopless, or M5 Ndc80 was essentially indistinguishable in the absence (**Figure 7B**, conditions 1-3) or presence (conditions 4-6) of the antibody. This result indicates that AB-850 does not interfere with clustering of the wild-type Ndc80 complex in cells, although we cannot exclude that this is because the complex dissociates after electroporation (as clarified above, the loopless and M5 mutants were not expected to be recognized efficiently by the AB-850). Cells electroporated with wild-type Ndc80 and AB-849 failed to congress their chromosomes, activated the SAC, but exited mitosis after arresting for ∼5 hours (**Figure 7B**, condition 7). Thus, crosslinking of Ndc80 complexes with AB-849 prevented proper kinetochore-microtubule interactions. A titration of AB-849:Ndc80 ratios revealed dose-dependent antibody effects on mitotic alignment, but not on the duration of cell division (**Suppl. Figure 8B**).

**Figure 8.**
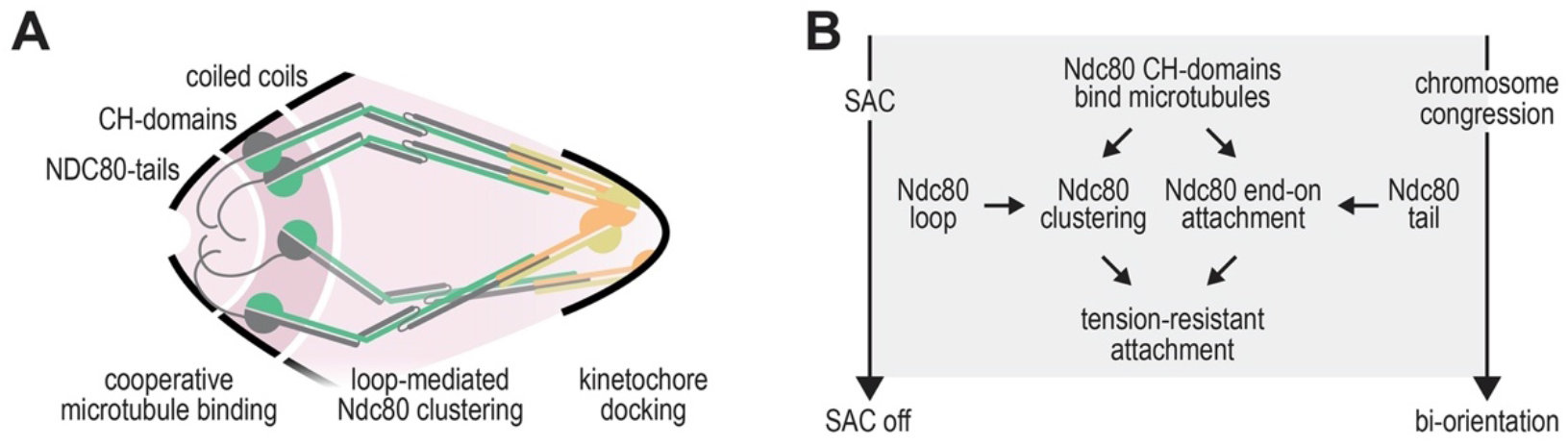
Synergistic contributions the Ndc80 loop and tail to robust kinetochore-microtubule binding. **A)** Schematic representation of four Ndc80 complexes. **B)** A model for the synergistic contributions of the Ndc80 loop and the Ndc80 tail to kinetochore-microtubule attachment.

When the mutated Ndc80 complexes (loopless and M5) were incubated with the AB-849 before electroporation, the electroporated cells exited mitosis rapidly (**Figure 7B**, conditions 8 and 9). This contrasts with the behavior observed in control cells receiving these mutants (**Figure 7B**, conditions 2 and 3), which arrested robustly in mitosis (see also **Figure 4**). This effect was not caused by restoration of chromosome congression (**Suppl. Figure 8**). Interestingly, although cells with wild-type and mutated Ndc80 loops both fail to congress their chromosomes properly in the presence of AB-849, cells with wild-type Ndc80 loops were able to maintain a SAC response whereas cells with the loopless and M5 mutants failed to arrest (**Figure 7B**, conditions 7-9). Collectively, these results imply that binding of the AB-849 antibody weakens SAC signaling and that this effect exposes a silent, underlying checkpoint defect caused by the loop mutants.

To test this idea further, we first monitored the duration of the SAC arrest in nocodazole-treated cells depleted of endogenous Ndc80 and expressing wild-type, loopless, or M5 Ndc80 (**Figure 7B**, conditions 10-12). All three complexes maintained a robust arrest. Co-electroporation with AB-849, on the other hand, dramatically decreased the duration of the arrest in presence of the loopless and M5 mutants, whereas it weakened the SAC only mildly in the presence of the wild-type Ndc80 (**Figure 7B**, conditions 13-15). These observations indicate that the region immediately preceding the loop (recognized by the AB-849) and the loop itself are important for SAC signaling. While individual perturbations at these sites are compatible with SAC signaling, concomitant perturbations result in a profound weakening of the SAC response.

## DISCUSSION

The loop region of the NDC80 subunit was previously identified for its essential role in the establishment of load-bearing interactions between kinetochores and microtubule ends required for chromosome congression (Maure et al., 2011; Shrestha and Draviam, 2013). Our finding in a minimally reconstituted system that the loop is dispensable for microtubule end-tracking in the absence of force, but crucial to stall and rescue microtubule shortening under force, supports this view (compare **Figure 2H,I** and **Figure 3B**). At a mechanistic level, however, the function of the loop has remained elusive and partly controversial, as discussed below. We demonstrate here that the NDC80 loop promotes Ndc80-Ndc80 interactions that are crucial to generate force-resistant attachments of modules with multiple immobilized Ndc80 complexes. Our observations also suggest that interactions between adjacent Ndc80 complexes may signal the establishment of load-bearing kinetochore-microtubule attachments and silence the SAC (**Figure 8**). We have not yet demonstrated a direct molecular link between Ndc80 clustering mediated by the loop and SAC signaling. We hypothesize the existence of a binding site for a SAC protein in a region encompassing the epitope recognized by the AB-849 antibody and the neighboring loop. In this model, presence of the antibody or deletion of the loop would, in isolation, cause an only partial loss of the SAC protein, while their combination would ablate recruitment and cause a SAC failure.

Loop-dependent binding of adjacent Ndc80 complexes requires binding to microtubules: full-length Ndc80 complexes are stable monomeric complexes in solution at concentrations as high as 50 μM (∼10 mg/ml) but cluster on microtubules at concentrations as low as 0.005 μM (**Figure 1**). This likely reflects that a microtubule orients and concentrates Ndc80 complexes in a manner that is sufficient to trigger homotypic Ndc80 interactions that require the loop. Although our structural analysis provides insight into the folding of the Ndc80 loop, a molecular explanation for loop-dependent clustering of Ndc80 complexes on microtubules is lacking. Since point mutations phenocopy deletion of the loop, both in dividing cells and in a reconstituted system, it appears likely that the loop directly binds an adjacent Ndc80 complex at a hitherto unidentified site.

The unstructured N-terminal tail of the NDC80 subunit, encompassing the protein’s first 80 residues, plays an important role in the coordinated binding of Ndc80 to the ends of dynamic microtubules *in vivo* and *in vitro* (Cheeseman et al., 2006; DeLuca et al., 2006; Guimaraes et al., 2008; Miller et al., 2008; Helgeson et al., 2018; Huis in ‘t Veld et al., 2019; Wimbish et al., 2020; Shrestha et al., 2017; DeLuca et al., 2018; Kucharski et al., 2022; DeLuca et al., 2011; Zhu et al., 2013; Long et al., 2017). We suggest that the Ndc80 tail and loop contribute to the establishment of tension-resistant kinetochore-microtubule attachments in a synergistic manner: the tails by coordinating the direct binding of Ndc80 complexes to the end of microtubules and the loops by stabilizing Ndc80 clusters at a distance from the microtubule-binding site (**Figure 8A**).

Cooperative microtubule binding of Ndc80 assisted by the Ndc80 tail has been demonstrated previously (Ciferri et al., 2008; Alushin et al., 2010, 2012; Janczyk et al., 2017). Notably, these studies were performed with a truncated version of the Ndc80 complex, Ndc80-bonsai, that lacks the loop and almost the entire coiled-coil stalk of the complex, and using Ndc80-bonsai concentrations in the low μM range. In contrast, we observe clustering on microtubules of full-length Ndc80 complexes at low nM concentrations. At similarly low concentrations, Ndc80-bonsai (with or without mutations in the tail) binds microtubules in a non-cooperative manner (Zaytsev et al., 2015).

Collectively, our findings allow us to formulate a new hypothesis on the coordination of the molecular events that mark the process of bi-orientation. We surmise that force-resistant microtubule binding by Ndc80 complexes, at its kinetochore concentration or at a realistically low concentration *in vitro*, requires i) the deployment of Ndc80 loops to trigger local clustering and ii) the deployment of Ndc80 tails at the end of the microtubule, facilitated by their dephosphorylation (**Figure 8B**). This model predicts that the Ndc80 complexes are oriented as parallel “pillars” and is therefore compatible with the high nematic order observed for Ndc80 *in vivo* (Roscioli et al., 2020). The model may also explain why even a non-phosphorylatable version of the Ndc80 tail cannot rescue the dramatic effects on bi-orientation caused by deletion of the loop. The N-terminal tails may be insufficient for clustering and coordinated microtubule binding when the loop is absent. A question for future work is how many adjacent CH-domains and tails are required to generate a robust end-on microtubule attachment. Furthermore, our AF2 predictions and structural analyses suggest that additional conserved elements of the Ndc80 complex may contribute to successful bi-orientation. Most notably, and as previously observed in micrographs and in predictions, there is a prominent disruption of the coiled coil in between the microtubule-binding region and the loop of Ndc80 (**Figure 1B-C**) (Jenni and Harrison, 2018). We propose to name this conserved element, encompassing NDC80^359-362^ and NUF2^244-247^ in humans, the Ndc80 hinge. How the hinge contributes to Ndc80-microtubule binding and chromosome biorientation remains to be addressed. Our analysis, however, indicates that the hinge, not the loop, is the likely primary site of bending in the first half of the Ndc80 shaft.

Importantly, the establishment of force-bearing kinetochore-microtubule attachments *in vivo* requires, in addition to the Ndc80 complex, additional microtubule binders, including the SKA and SKAP/Astrin complexes (Monda and Cheeseman, 2018). It is therefore possible that Ndc80 multivalency, in addition to having a direct effect on microtubule binding, also controls the interaction with these additional linkages. Indeed, a crucial role of the loop in kinetochore-microtubule attachment was previously demonstrated in a range of organisms (Hsu and Toda, 2011; Maure et al., 2011; Varma et al., 2012; Zhang et al., 2012; Tang et al., 2013; Shrestha and Draviam, 2013; Scarborough et al., 2019; Wimbish et al., 2020). These studies postulated that the loop promotes stable kinetochore-microtubule attachment through the direct binding of microtubule binding proteins such as the Ska complex, CH-TOG/Stu2, or Cdt1 (Hsu and Toda, 2011; Varma et al., 2012; Zhang et al., 2012; Tang et al., 2013; Zhang et al., 2017). Subsequent studies, however, questioned the notion that the loop is directly involved in these physical interactions. For instance, recruitment of Dam1 to Ndc80 complexes in budding yeast appears to require the Ndc80 loop *in vivo*, but not *in vitro* (Maure et al., 2011; Lampert et al., 2013; Jenni and Harrison, 2018). Similarly, the recruitment of the Ska complex to kinetochores requires the Ndc80 loop and the Ndc80 tail *in vivo* (Zhang et al., 2012; Janczyk et al., 2017), but Ska and Ndc80 form a stable stoichiometric complex *in vitro* in a loop- and tail-independent manner, at least at micromolar concentration (Huis in ‘t Veld et al., 2019). Thus, lack of recruitment of certain downstream proteins to kinetochores with mutated Ndc80 loops might not reflect a direct role of the loop in these recruitments, but rather that the Ndc80 clusters assembled through the loop are stronger binding platforms for these proteins than the isolated Ndc80 complexes. This possibly reflects preferred interactions of Ndc80 clusters on microtubules with other multimers, such as the Ska and Dam1 complexes. In the future, we will ascertain the validity of this hypothesis and test why the late recruitment of certain protein to the kinetochore requires an intact loop.

Mutation of the loop increases the diffusion of Ndc80 on microtubules. Interactions between loopless Ndc80 complexes, induced with an antibody that binds near the loop, reduced the diffusion of loopless Ndc80 trimers (**Figure 6**). This demonstrates that the binding between adjacent Ndc80 complexes increases the grip to the microtubule lattice. Disturbing the diffusion of Ndc80 ensembles on microtubules might hinder proper chromosome congression and end-on microtubule binding in cells. Consistently, the electroporation of antibody:Ndc80 mixtures into cells interfered with chromosome congression and checkpoint signalling (**Figure 7B**). These results are consistent with classic studies demonstrating that i) antibodies against NDC80 or NUF2 injected into Xenopus cells resulted in a mitotic arrest and chromosome congression failure (McCleland et al., 2003) and ii) a monoclonal antibody (9G3, raised against GST-NDC80^56-642^, later shown to recognize NDC80^200-215^) interfered with mitosis by hyperstabilizing kinetochore-microtubule connections (Chen et al., 1997; DeLuca et al., 2006). Interestingly, we found that the 9G3 antibody binds microtubules *in vitro*, preventing the analysis of its influence on the diffusion of Ndc80 trimers (**Suppl. Figure 7**).

The kinetochore-microtubule interface consists of dozens of proteins and protein complexes, all present in multiple copies. Understanding how these proteins control the attachments of chromosomes to the mitotic spindle and link this attachment state to cell cycle progression is a formidable challenge. In this study, we investigated how a short sequence in the outer kinetochore, the Ndc80 loop, coordinates the formation of stable end-on kinetochore-microtubule attachments that support chromosome congression and biorientation. To unravel the multifaceted roles of the loop, we identified and tested separation-of-function mutants under the controlled conditions of a fully reconstituted system and in the presence of additional proteins and regulatory pathways in cells. In the future, our approach can provide molecular and mechanistic insight into other central aspects of kinetochore biology, such as the transition from a lateral to an end-on attachment, the role that other microtubule binders play, and how intricate feedback loops control kinetochore-microtubule binding.

## Supporting information

Supplementary data iii

## AUTOR CONTRIBUTIONS

### Conceptualisation

SP, AM, VAV, PJH; Methodology, Data Curation, Software: SP, IRV, VAV, PJH; Investigation, Formal analysis, Validation: SP, HM, IRV, VAV, PJH; Resources: MT, ADA; Supervision, Project administration: SP, MD, AM, VAV, PJH; Funding acquisition: MD, AM; Visualisation: VAV, PJH; Writing – original draft preparation: AM, VAV, PJH; Writing – review and editing: all authors.

## ACKNOWLEDGEMENTS

We thank Christina Courtis, Alicia Dammers, Farzad Khanipour, Laura Ney, and Isabelle Stender for help with the generation of recombinant Ndc80 complexes. We are grateful to Petra Geue and Raphael Gasper-Schoenenbruecher for assistance with SEC-MALS and mass photometry, and to Franziska Müller, Andreas Brockmeyer, Malte Metz, and Petra Janning for mass spectrometry. VAV was supported by the European Research Council Synergy Grant 609822 to MD, and by the QMUL Startup grant (SBC8VOL2). AM acknowledges funding of this work through the European Research Council Synergy Grant SyG 951430 BIOMECANET.

## SUPPLEMENTARY DATA

i. A PDB file of the predicted structure of the full-length human Ndc80 complex. Corresponding to **Figure 1** and **Suppl. Figure 1**.
ii. A Chimera X scene with the predicted structures of full-length Ndc80 complex from human and budding yeast and the predicted structures of the loop region in various organisms (aligned to the human fragment NDC80^376-516^:NUF2^269-356^). Corresponding to **Suppl. Figure 2**. Please email, .cxs not accepted by bioRxiv.
iii. A table containing all unique intra- and intermolecular Ndc80 crosslinks identified with a false-discovery rate of 1% Crosslinking-ms table. Corresponding to **Suppl. Figure 4**.

## MATERIALS and METHODS

### Molecular Modeling

The version AF2 Multimer 2 (Evans et al., 2021) of AlphaFold 2 (AF2) was used for all molecular modeling. Compared to the original AF2 (Jumper et al., 2021), AF2 Multimer is more sensitive to intra- and intermolecular interactions, resulting in a tendency to collapse extended proteins, *e*.*g*. long coiled-coil regions, into more compact structures. To avoid this, sub-fragments with overlapping regions were modeled separately and then stitched together. Fragment length was a compromise between “long and bending” and “short with dissociated coiled coils at the ends”. Several different regions were predicted and analysed, and we selected the longest possible fragments that did not bend. Three segments with overlapping regions were stitched together: i) NDC80 81-401 | NUF2 1-283, ii) NDC80 318-522 | NUF2 205-359, and iii) NDC80 507-642 | NUF2 344-464. Models were then superimposed on their overlapping parts using PyMOL (The PyMOL Molecular Graphics System, Version 2.5.2 Schrödinger, LLC.) and a residue in the middle of the best-fitting regions was chosen as fragment boundary. The final model was comprised of i) NDC80 81-382 | NUF2 1-267, ii) NDC80 383-511 | NUF2 268-351, and iii) NDC80 512-642 | NUF2 352-464 (**Suppl. Figure 1**). The ends of the fragments tend to have higher pLDDT scores, which is expected due to the truncated multiple sequence alignment that is then used by AF2. The geometry at the stitching points was checked and optimized with COOT (Emsley and Cowtan, 2004), followed by a minimisation of the complete model with PHENIX (Liebschner et al., 2019). Backbone restraints were applied to conserve the AF2-predicted backbone positions. In the final model, 98% of the residues had pLDDT scores above 80 (high confidence) and 84% of the residues had scores above 90 (very high confidence) (**Suppl. Figure 1**). Corroborating the high quality of the model, the predicted alignment error (PAE) plots consistently indicated well-defined relative positions of the monomers, including the tetramerisation domain that is composed of SPC24, SPC25, and the C-termini of NDC80 and NUF2 (**Suppl. Figure 1**). Omitting the structural template for the yeast tetramerisation domain (PDB ID 5TCS) (Valverde et al., 2016) from the AF2 database did not change the prediction for the human tetramerisation domain. The loop region of NDC80 (residues 422-458) was predicted very reproducibly, whereas the angle of the hinge (residues 358-363 in NDC80 and 243-248 in NUF2) varied between 109° to 131° in various predictions. In addition, some predictions showed up to 15° lateral deviation of the coiled-coil axes perpendicular to the plane of the triangle. It is unclear if this is a *bona fide* property of the Ndc80 complex, but it is interesting that AF2 reproduces the observation of the hinge-like region of the Ndc80 at a distance of approximately 180-210 Å (+/-30 residues) from the globular, N-terminal NDC80/NUF2 domains (Huis in ‘t Veld et al., 2016; Jenni and Harrison, 2018), which corresponds well with the position of the predicted kink residues (358-363 in NDC80 and 243-248 in NUF2) in our model (approximate distance from N-terminus 170-200 Å).

To compare the evolutionary conservation of the loop region, NDC80 sequences from 33 diverse eukaryotic organisms were selected (Boeckmann et al., 2015) and aligned and visualized using proviz (Jehl et al., 2016) and jalview (Waterhouse et al., 2009) (**Suppl. Figure 2**). The phylogenetic tree was generated using ITOL (Letunic and Bork, 2021) and structures were predicted using AF2 as described above (**Suppl. Figure 2**). All figures were made using ChimeraX (v1.2 and daily builds) (Goddard et al., 2018).

### Cloning, expression, and purification of Ndc80

Expression cassettes from pLIB vectors containing NDC80, NUF2, SPC25^SORT-HIS^, and SPC24 were combined on a pBIG1 vector using Gibson assembly as described (Weissmann et al., 2016; Volkov et al., 2018). Baculoviruses were generated in *Sf9* insect cells and used for protein expression in *Tnao38* insect cells. Between 60 and 72 hours post-infection, cells were washed in PBS and stored at -80 °C. All subsequent steps were performed on ice or at 4 °C. Cells were thawed and resuspended in lysis buffer (50 mM Hepes, pH 8.0, 200 or 250 mM NaCl, 10% v/v glycerol, 2 mM TCEP, 20 mM imidazole, 0.5 mM PMSF, protease-inhibitor mix HP Plus (Serva)), lysed by sonication and cleared by centrifugation at 108,000g for 60 minutes. The cleared lysate was filtered (0.8 μM) and applied to a 10 or 20 ml HisTrap FF (GE Healthcare) equilibrated in washing buffer (lysis buffer without protease inhibitors). The column was washed with approximately 20 column volumes of washing buffer and bound proteins were eluted with elution buffer (washing buffer containing 300 mM imidazole). Relevant fractions were pooled, diluted 5-fold with buffer A (50 mM Hepes, pH 8.0, 25 mM NaCl, 5% v/v glycerol, 1 mM EDTA, 2 mM TCEP) and applied to a 25 ml Source15Q (GE Healthcare) strong anion exchange column equilibrated in buffer A. After washing with approximately 20 column volumes, bound proteins were eluted with a linear gradient from 25 mM to 300 mM NaCl in 180 ml. Relevant fractions were concentrated in 50 kDa molecular mass cut-off Amicon concentrators (Millipore). Complexes were then fluorescently labelled (see next section) or directly applied to a Superdex 200 16/600 increase, a Superose 6 10/300 increase, or a Superose 6 prep grade XK 16/600 column (GE Healthcare). Columns were equilibrated in Ndc80 buffer (50 mM Hepes, pH 8.0, 250 mM NaCl, 5% v/v glycerol, 2 mM TCEP) and size-exclusion chromatography was performed under isocratic conditions at recommended flow rates. Relevant fractions were pooled, concentrated, flash-frozen in liquid nitrogen, and stored at -80 °C.

### Fluorescent labelling of Ndc80

The calcium independent Sortase 7M (Hirakawa et al., 2015) was used for the C-terminal conjugation of a synthetic peptide to SPC25. Various synthetic peptides (Genscript) were used for this purpose: GGGGK^FAM^, GGGGK^TMR^, and GGGGC^Alexa-488^. Reactions were performed at 10 °C for 4-16 hours in Ndc80 buffer with Sortase:Ndc80:Peptide ratios of approximately 1:5:25 with Ndc80 in the 10-20 μM range. Fluorescently labelled complexes were purified using size-exclusion chromatography, as described above.

### Assembly of Ndc80 trimers

Ndc80 complexes were coupled to streptavidin-derived T_1_S_3_ scaffolds as described (Volkov et al., 2018) (**Suppl. Figure 3**). In brief, T1S3 scaffolds were assembled from core traptavidin (T; addgene plasmid #26054) and Dead Streptavidin-SpyCatcher (S; addgene plasmid # 59547) (Chivers et al., 2010; Fairhead et al., 2014). T_1_S_3_ scaffolds were incubated with an approximate 10-fold molar excess of Ndc80 for 12-20 hours at 10 °C in the presence of PMSF (1 mM) and protease inhibitor mix (Serva). Sortase labeling was achieved in the same reaction, as described above. Reaction mixtures were applied to a Superose 6 increase 10/300 column (GE Healthcare) equilibrated in 20 mM TRIS pH 8.0, 200 mM NaCl, 2% v/v glycerol, 2 mM TCEP. Size-exclusion chromatography was performed at 4 °C under isocratic conditions at recommended flow rates and the relevant fractions were pooled and concentrated using 30 kDa molecular mass cut-off Amicon concentrators (Millipore), flash-frozen in liquid nitrogen, and stored at -80 °C.

### SEC-MALS and mass photometry

Full-length and loopless Ndc80 complexes were analysed by SEC-MALS on a Dawn Heleos II System with an Optilab T-rEX RI detector (Wyatt) and a 1260 Inifinity II LC system (Agilent). The Superose 6 increase 10/300 column (GE Healthcare) was pre-equilibrated with glycerol-free Ndc80 buffer (50 mM HEPES pH 8.0, 250 mM NaCl, and 2mM TCEP). Analysis was performed at room temperature with 60 μl full-length or loopless Ndc80 complex that was diluted in running buffer to 2 mg/ml. Mass photometry was performed on a Refeyn Two^MP^ System (Refeyn) that was calibrated with a mixture of BSA (66.5 and 123 kDa) and Thyroglobulin (330 and 660 kDa). Ndc80 complexes were diluted to a concentration of 100 nM in glycerol-free Ndc80 buffer and analysed in 20 μL droplets following a ten-fold dilution in buffer to 10 nM. Data were analysed and plotted using the Discover^MP^ software (Refeyn).

### Crosslinking and mass spectrometry

200 μL of full-length Ndc80, loopless Ndc80, and their complexes with the Mis12 complex were prepared at 7.5 μM in crosslinking buffer (50 mM HEPES pH 8, 250 mM NaCl, 5% v/v glycerol, and 2 mM TCEP), crosslinked with an approximate 500-fold molar excess of DSBU (3.75 mM), and quenched with TRIS (100 mM) as described in **Suppl. Figure 4**. Samples were prepared for mass spectrometry and analysed using MeroX (Version 2.0.1.4) as previously described (Arlt et al., 2016; Pan et al., 2018). Crosslinks were inspected in XiView (Graham et al., 2019) and, after setting a false-discovery rate to 1%, ranked according to the number of matched peptides and the maximum score. Unique intra- and intermolecular Ndc80 crosslinks from all four datasets were identified. To distinguish frequently observed and more rarely observed crosslinked peptides, the rank was used to divide the dataset, at an arbitrary cut-off, in two parts. All crosslinks were displayed on the predicted structure of full-length Ndc80 and C_α_-C_α_ distances were plotted using ChimeraX (daily build) (Goddard et al., 2018).

### Low-angle metal shadowing and electron microscopy

Full-length and loopless Ndc80 complexes (10 μM) were incubated with full-length Mis12 complexes (15 μM) for approximately 1 hour on ice. Mixtures were diluted 1:9 with Ndc80 buffer, diluted 1:1 with spraying buffer (200 mM ammonium acetate and 60% glycerol), and then air-sprayed onto freshly cleaved mica (V1 quality, Plano GmbH) of approximately 2×3 mm. Specimens were mounted and dried in a MED020 high-vacuum metal coater (Bal-tec). A Platinum layer of approximately 1 nm and a 7 nm Carbon support layer were evaporated subsequently onto the rotating specimen at angles of 6-7° and 45° respectively. Pt/C replicas were released from the mica on water, captured by freshly glow-discharged 400-mesh Pd/Cu grids (Plano GmbH), and visualized using a LaB6 equipped JEM-400 transmission electron microscope (JEOL) operated at 120 kV. Images were recorded at a nominal magnification of 60,000x on a 4k X 4k CCD camera F416 (TVIPS), resulting in 0.1890 nm per pixel. Particles were manually selected using EMAN2 (Tang et al., 2007).

### Tubulin and microtubules

Pig brain tubulin, as well as TMR- and HiLyte488-labelled tubulin, were purchased from a commercial source (Cytoskeleton Inc). DIG-tubulin was made in-house by labelling microtubules polymerized from commercial tubulin with an NHS-ester (Sigma Aldrich) of digoxigenin according to published protocols (Hyman et al., 1991). GMPCPP-stabilized seeds were made by two rounds of polymerisation at 37°C in presence of 1 mM GMPCPP and 25 μM tubulin (40% DIG-labelled). Seeds were sedimented in a Beckman Airfuge at 126,000g for 5 min. After the second polymerisation round, the seeds were resuspended in MRB80 buffer (80 mM K-Pipes pH 6.9, 1 mM EGTA, 4 mM MgCl^2^) supplemented with 10% glycerol and aliquots were flash-frozen in LN_2_.

To prepare taxol-stabilized microtubules, 70 μM tubulin (with or without DIG- and fluorescent labels) was polymerized in presence of 25% glycerol and 1 mM GTP for 10 min at 37°C, and then stabilized by an addition of 25 μM taxol followed by a 20 min incubation. Microtubules were sedimented in a Beckman Airfuge at 32,000g for 3 min, resuspended in MRB80 buffer supplemented with 40 μM taxol and stored at 25°C for up to three days.

### Preparation of flow chambers and TIRF microscopy

Microscopy chambers were prepared using slides and coverslips treated with oxygen plasma and immediately silanized with a mixture of 2% dichlorodimethyl silane and 98% octamethylcyclooctasilane) for 5 min (Gell et al., 2010; Maleki et al., 2022). Before an experiment, antibodies against DIG (Roche 11333089001) or tubulin (TU-20, Abcam) were adsorbed to the silanized glass, followed by a passivation with 1% Pluronic F-127 in MRB80. Taxol-stabilized microtubules were diluted in approximately 5-7 chamber volumes of MRB80 with 10 μM taxol, washed in within 3 min and immediately washed out with three chamber volumes of MRB80 with 10 μM taxol to produce microtubules mostly oriented along the chamber. Subsequently, proteins of interest were introduced into the chamber in imaging buffer containing MRB80, 0.1% methylcellulose, 40 μM taxol, 1 mg/ml κ-casein, 4 mM DTT, 0.2 mg/ml catalase, 0.4 mg/ml glucose oxidase and 20 mM glucose. Imaging buffer was pre-cleared in a Beckman Airfuge for 5 min at 126,000g before the addition of Ndc80 or antibodies.

For experiments with dynamic microtubules, DIG-labelled GMPCPP-stabilized microtubule seeds were attached to silanized chambers containing anti-DIG antibodies and passivated with Pluronic F-127. Imaging buffer was prepared as described above, however taxol was substituted with 11 μM tubulin (3-5% labelled).

Experiments involving monomeric Ndc80 were performed using a Nikon Eclipse Ti2 microscope equipped with a Plan Apo 100 × 1.45 NA TIRF, iLas^3^ ring TIRF illumination system (GATACA), and an Andor iXon 897 EMCCD camera. All other experiments were performed with a Nikon Ti-E microscope equipped with a Plan Apo 100 × 1.45 NA TIRF oil-immersion objective, iLas^2^ ring TIRF module (Roper Scientific) and a Evolve 512 EMCCD camera (Roper Scientific). Images were recorded using MetaMorph software. Experiments with taxol-stabilized microtubules were performed at ambient temperature (23°C); for experiments with dynamic microtubules the objective was heated to 34°C using a custom-made collar coupled with a thermostat, resulting in the flow chamber being heated to 30°C.

### Image analysis

Coordinates and brightness of particles diffusing on microtubule lattice or tracking their ends were analysed using Fiji (Schindelin et al., 2012) and Julia using custom scripts available at https://github.com/volkovdelft/kymo.jl. Kymographs were made through a reslice operation using the kymograph_(n)channel.ijm macro. Position of a particle was determined in each line of the kymograph using kymo.ipynb jupyter notebook and following in-line comments. Oligomerisation was inferred from initial brightness of a particle and the photobleaching step characteristic for a fluorophore as described previously (Volkov et al., 2018). Mean squared displacement curves to characterize one-dimensional diffusion were generated using process_diffusion.ipynb: coordinates of each diffusing spot were iteratively split into segments of equal length with an offset of one datapoint, and subsequently aligned at their start and averaged. To analyze co-localisation of Ndc80 monomers and trimers on microtubules, trimers were tracked in one fluorescent channel of a kymograph as above using kymo_2channel.ipynb, and then the brightness of monomers was measured in the second fluorescent channel at the coordinates determined for trimer. The resulting monomer brightness was thresholded using a value for single Alexa488 fluorophore, and the number of datapoints above the threshold was divided by the total number of datapoints to calculate the time fractions for co-localisation.

### Preparation of beads and force measurements

Beads were prepared as described previously (Volkov et al., 2018). In brief, 1 μm glass COOH-functionalized beads were coated covalently with PLL-PEG (Poly-L-lysine (20 kDa) grafted with polyethyleneglycole (2 kDa), SuSoS AG, Switzerland) containing varying fraction of biotinylated PLL-PEG of the same composition. Biotins on the beads were then saturated with streptavidin-oligomerized Ndc80-TMR trimers. Bead preparations were analysed for bead brightness using at least 50 single beads.

Optical trapping was performed using a custom optical setup as described previously (Volkov et al., 2018). A bead coated with Ndc80 trimers was captured in a trap with a stiffness of 0.2-0.4 pN/nm placed near a growing end of a dynamic microtubule. Bead-microtubule attachment was verified by switching the trap off for several seconds and monitoring bead’s motion: successfully attached beads moved in an arc shape across the direction of microtubule growth and did not move significantly along it. The bead was then re-trapped again and the experiment was initiated by simultaneous start of acquisition of DIC images of the bead-microtubule pair, and the quadrant photo detector (QPD) readings to monitor bead’s displacement from the trap center. Recording was stopped after the microtubule depolymerized and the bead detached from it, or at will after 30-40 min of data acquisition if the bead did not detach. In the latter case, the trap stiffness was increased 10 times, and the bead was ruptured from the microtubule using 100 nm steps of the piezo stage. Motions of a free bead were recorded after detachment for each bead to later use them to determine trap stiffness.

### Cell electroporation

All electroporation experiments of this study were performed as described using a Neon Transfection System Kit (Thermo Fisher) (Alex et al., 2019). Cells were collected from the surface of the cell plate by trypsinisation, washed two times with PBS, and counted. Between 2 to 3 million cells were used per electroporation and were resuspended in 90 μl of buffer R (Thermo Fisher). Proteins were spun down at 16700 *g* for 10 minutes and then diluted with buffer R in a 1:1 ratio. For electroporation with antibody, cells were resuspended in 56 μl of buffer R and the Ndc80-antibody mixture was diluted 1:1 in buffer R. The Ndc80-antibody and cell suspension were mixed and then the mixture was loaded into a 100 μl Neon Pipette Tip (Thermo Fisher). Electroporation was performed at 1005 V with two consecutive pulses of 35 msec. Cells were washed with PBS and then treated with Trypsin to digest non-cell-internalized proteins. After another wash with PBS, cells were plated on 6-well plates for immunofluorescence staining or 24-well cell-imaging plates (Ibidi) for live-cell imaging.

### Antibody generation

Peptides *C-*TQLAEYHKLARKLKLI PKGAENS*-NH*_*2*_ (NDC80^410-432^), and *Ac-*AENSKGYDFEIKFNPEAGANCL*-NH*_*2*_ (NDC80^429-450^) were used to generate the antibodies AB-849 and AB-850, respectively. The peptides, synthesized at a minimal purity of 80% and conjugated to Keyhole Limpet Hemocyanin carrier (Mimitopes), were used for rabbit immunization (Eurogentec). Polyclonal antibodies were affinity purified from sera using peptides (elution pH 2.5) and stored at 4°C in PBS supplemented with 150 mM KCl, 0.1% BSA, and 0.05% NaN_3_.

### Cell culture, siRNA transfection, immunoblotting

HeLa, mCherry-H2B HeLa cell lines were cultured in DMEM (PAM Biotech) supplemented with 10% FBS (Clontech), 2 mM L-glutamine (PAN Biotech), 1% Penicillin, Streptomycin (Gibco). Cells were grown in a humidified chamber at 37°C in the presence of 5% of CO_2_. Cell lines were checked regularly for mycoplasma contamination and tested negative. Depletion of endogenous Ndc80 complex was attained by RNAiMAX (Invitrogen) transfection of siRNAs targeting three of the four subunits of the Ndc80 complex as described (Kim and Yu, 2015). The siRNA oligos were GAGUAGAACUAGAAUGUGA (siNdc80-4), GGACACGACAGUCACAAUC (siSpc24), and CUACAAGGAUUCCAUCAAA (siSpc25). To generate lysate of cells for immunoblotting, cells treated with control or Ndc80 complex siRNA were collected by trypsinisation and resuspended in lysis buffer [150 mM KCl, 75 mM Hepes pH 7.5, 1.5 mM EGTA, 1.5 mM MgCl_2_, 10 % Glycerol, and 0.075 % NP-40 supplemented with protease inhibitor cocktail (Serva)]. Lysates were run in 4-12 % gradient gels (NuPAGE, ThermoFisher) and proteins were transferred to a Nitrocellulose membrane for further analysis. The following antibodies were used for the immunoblot analysis of this study: anti-Hec1 (human Ndc80; mouse clone 9G3.23 GeneTex, Inc; 1:1000), anti-Vinculin (mouse monoclonal; clone hVIN-1; Sigma-Aldrich; 1:10000), AB-849 and AB-850 (rabbit polyclonal, in-house generated, 1:500).

### Cell treatment, microscopy, immunofluorescence detection, and live-cell imaging

Cells were imaged with a customized 3i Marianas system (Intelligent Imaging Innovations) equipped with an Axio Observer Z1 microscope chassis (Zeiss), a CSU-X1 confocal scanner unit (Yokogawa Electric Corporation), Plan-Apochromat 100x/1.4 NA objectives (Zeiss), and an Orca Flash 4.0 sCMOS Camera (Hamamatsu). Slide book software was used to acquire images as Z sections. Additional images were acquired using a DeltaVision Elite System (GE Healthcare) equipped with an IX-71 inverted microscope (Olympus), a UPlanFLN 40×1.3 NA objective (Olympus), and a pco.edge sCMOS camera (PCO-TECH Inc.). Live cell movies were taken as Z scans every 10 min interval for 16 hours. The softWoRx software was used for maximal intensity projection and analysis of the movies. For immunofluorescence sample preparation, cells were grown on poly-d-Lysine (Millipore) coated coverslips and permeabilized with a 0.5 % Triton-X in PHEM buffer. Cells were fixed with 4% paraformaldehyde (PFA) and then blocked for one hour with 5% boiled donkey serum. The following antibodies were used for immunostaining: anti-Hec1 (human Ndc80; mouse clone 9G3.23 GeneTex, Inc; 1:1000), anti-CENP-C (guineapig polyclonal, MBL, 1:500), anti-CREST/anti-centromere antibodies (human, Antibodies Inc, 1:1000), anti-tubulin (mouse monoclonal, Sigma, 1:5000), Knl1 (rabbit polyclonal, in house, 1:500), Nsl1 (mouse monoclonal, in house, 1:800), CENP-T (rabbit polyclonal, in house, 1:500), BUB1 (rabbit polyclonal, Abcam, 1:500), BubR1 (rabbit polyclonal, Abcam, 1:400). DNA was visualized by staining with 0.5 μg/ml DAPI (Serva) and coverslips were mounted to slide with Mowiol mounting media. All images were processed using Fiji. Quantifications of protein signals were measured using a custom-written script. All signals were normalized to either CREST or CENP-C signal. To calculate times in mitosis, 16 hr (960 min) long live-cell movies were analysed manually. Cells that entered mitosis within the first 500 minutes of the movie were accounted for. Cells already in mitosis at the start of the movie were not analysed. Mitotic duration was calculated from the time that cells condensed their chromosomes until they exited mitosis. For cold shock assay, cells were kept on ice and ice-cold media was added. Cells were kept in this condition for 14 mins and then fixed with 4 % PFA and proceeded to immunofluorescence staining as mentioned above. For treatment with drugs, hesperidin (500 nM), reversine (250 nM), and nocodazole (3.3 μM) were added 1 hour before starting the live-cell movies.

---

**Suppl. Figure 1.**
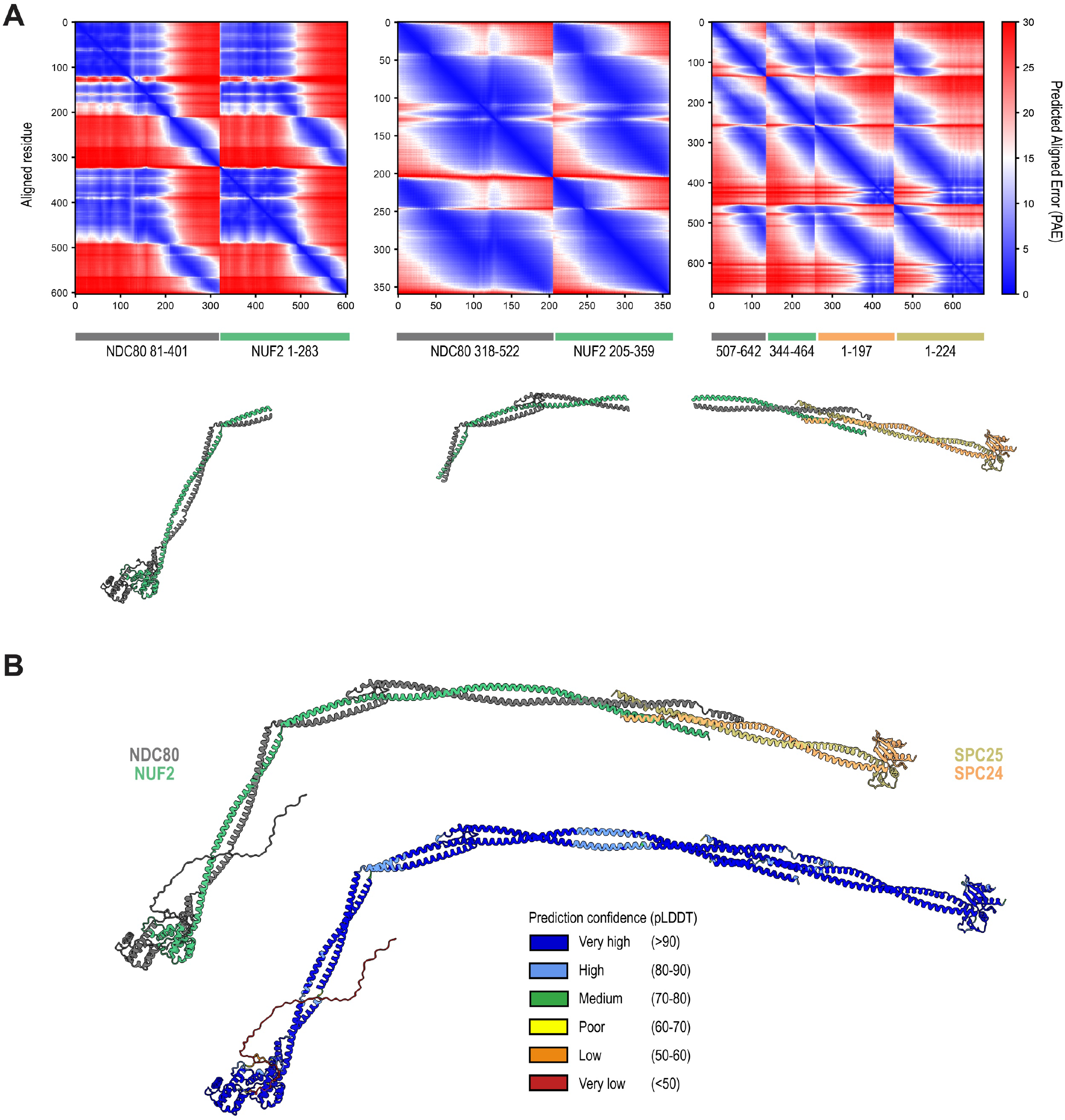
Structural *in silico* analysis of the human Ndc80 complex. **A**) Boundaries and Predicted Aligned Error (PAE) scores of the three Ndc80 segments that were predicted by AF2 multimer. These fragments were used to generate a composite prediction of the full-length Ndc80 complex. More information can be found in the Materials and Methods section. **B**) The prediction of the full-length Ndc80 complexes with colors representing the different subunits (as in **Figure 1B**) and the local prediction confidence intervals.

**Suppl. Figure 2.**
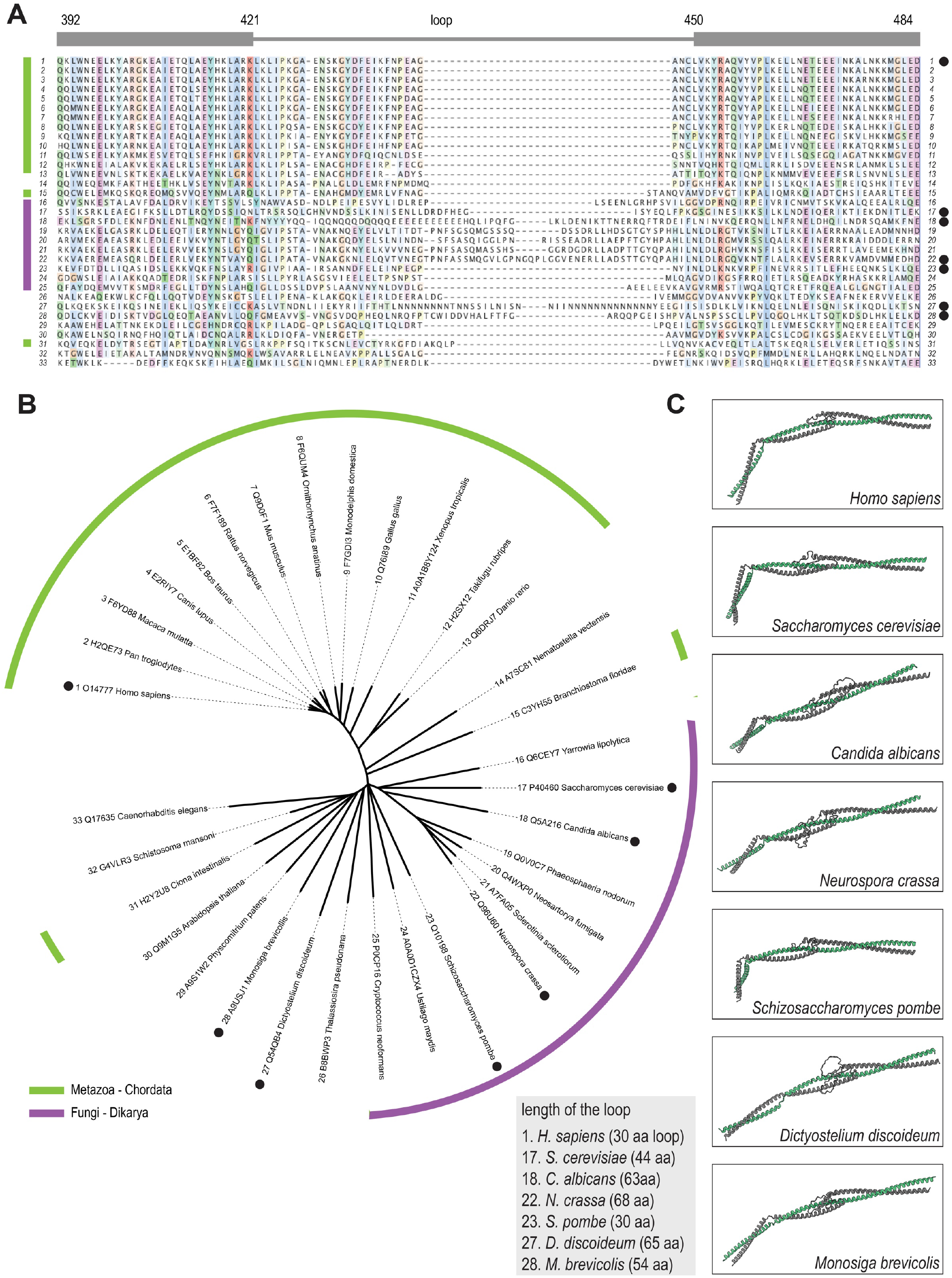
Alignments, phylogenetic tree, and structural conservation of the Ndc80 kink and loop. **A**) Sequence alignment of the loop region of the NDC80 subunit in various species. Residue numbers correspond to the human NDC80. **B**) Unrooted phylogenetic tree that was generated with complete NDC80 sequences. Sequences in panel A were arranged according to this tree. Species belonging to the Chordata phylum (light green) and the Fungi kingdom (purple) are indicated. Black dots mark species for which we predicted the structure. **C**) Predicted structures of the NDC80:NUF2 region spanning the hinge and loop regions. Shown in similar orientations following structural alignment to the human fragment NDC80^376-516^:NUF2^269-356^.

**Suppl. Figure 3.**
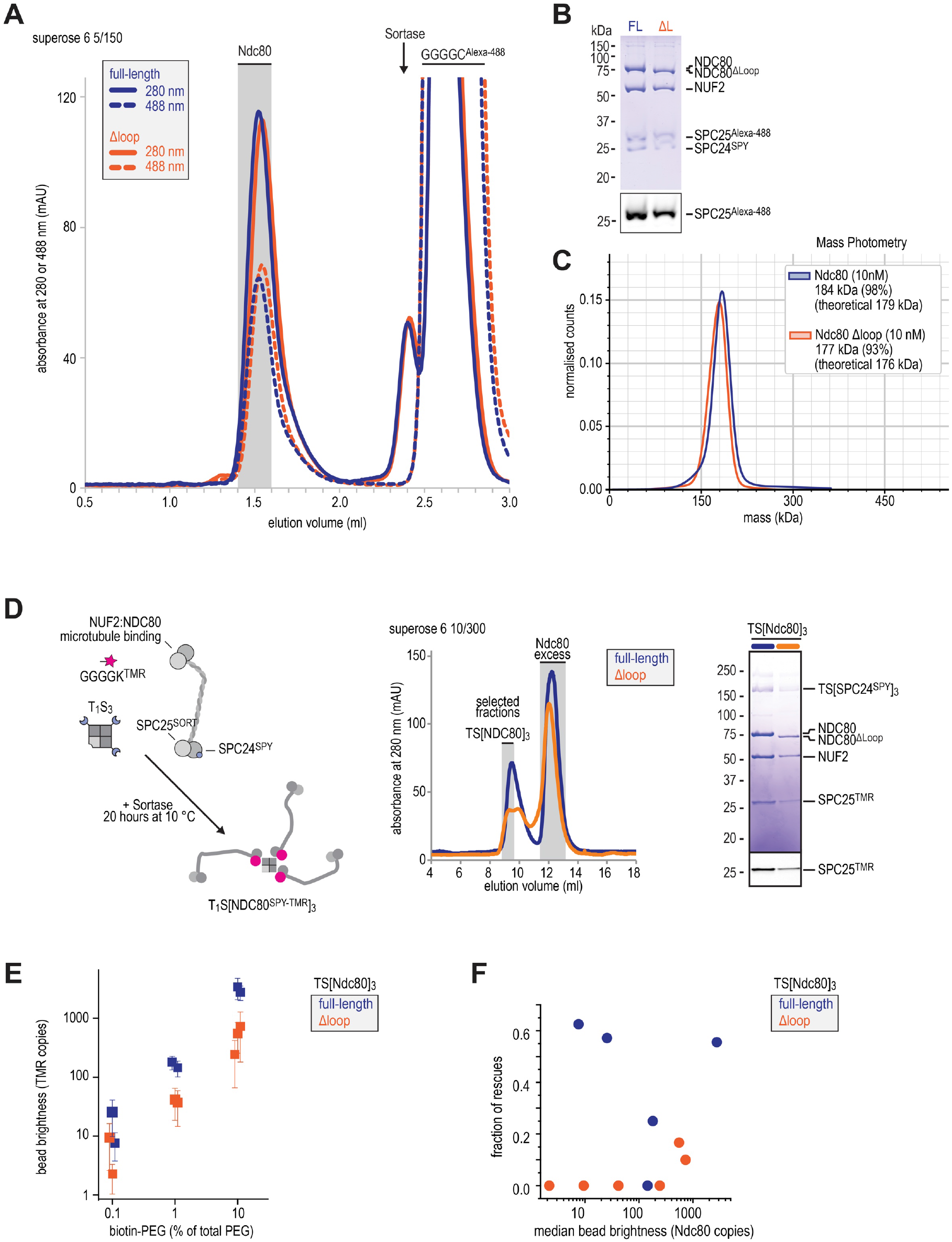
Preparation of full-length and loopless Ndc80, TS[Ndc80]_3_ modules, and coated beads. **A**) The fluorescent peptide, Sortase, and labelled Ndc80 complexes with (full-length, blue) and without (Δloop, orange) the loop were separated using size-exclusion chromatography. The gray area indicates Ndc80 that was collected and (without further concentration) stored for further use. **B**) Ndc80 complexes from panel A were analysed by in-gel fluorescence and Coomassie staining following SDS-PAGE. These complexes were used for experiments shown in **Figure 1F-K** and **Figure 5A-D. C**) Full-length and loopless Ndc80 complexes (Sortase labelled with FAM) were analysed by mass photometry. Determined and theoretical masses are indicated in the legend. These complexes were also used for the SEC-MALS shown in **Figure 1D. D**) Schematic overview of the preparation of Ndc80 trimers. The fluorescent peptide, Sortase, labelled Ndc80 monomers (full-length, blue; Δloop, orange), and Ndc80 trimers were separated using size-exclusion chromatography. Selected fractions containing Ndc80 trimers are marked in grey and were analysed by SDS-PAGE. Since samples were not boiled, the streptavidin scaffold and the covalently bound SPC24 subunits remain intact. See the Materials and Methods and (Volkov et al., 2018) for more information. **E**) Brightness of PLL-PEG-conjugated beads with various percentage of biotinylation, subsequently saturated with Ndc80^TMR^ trimers. Shown are mean and SD. Each datapoint represents a single bead preparation, at least 50 beads were quantified for each preparation. **F**) Fraction of stalls resulting in a rescue, binned by individual bead preparation, and correlated to the median bean brightness in that preparation

**Suppl. Figure 4.**
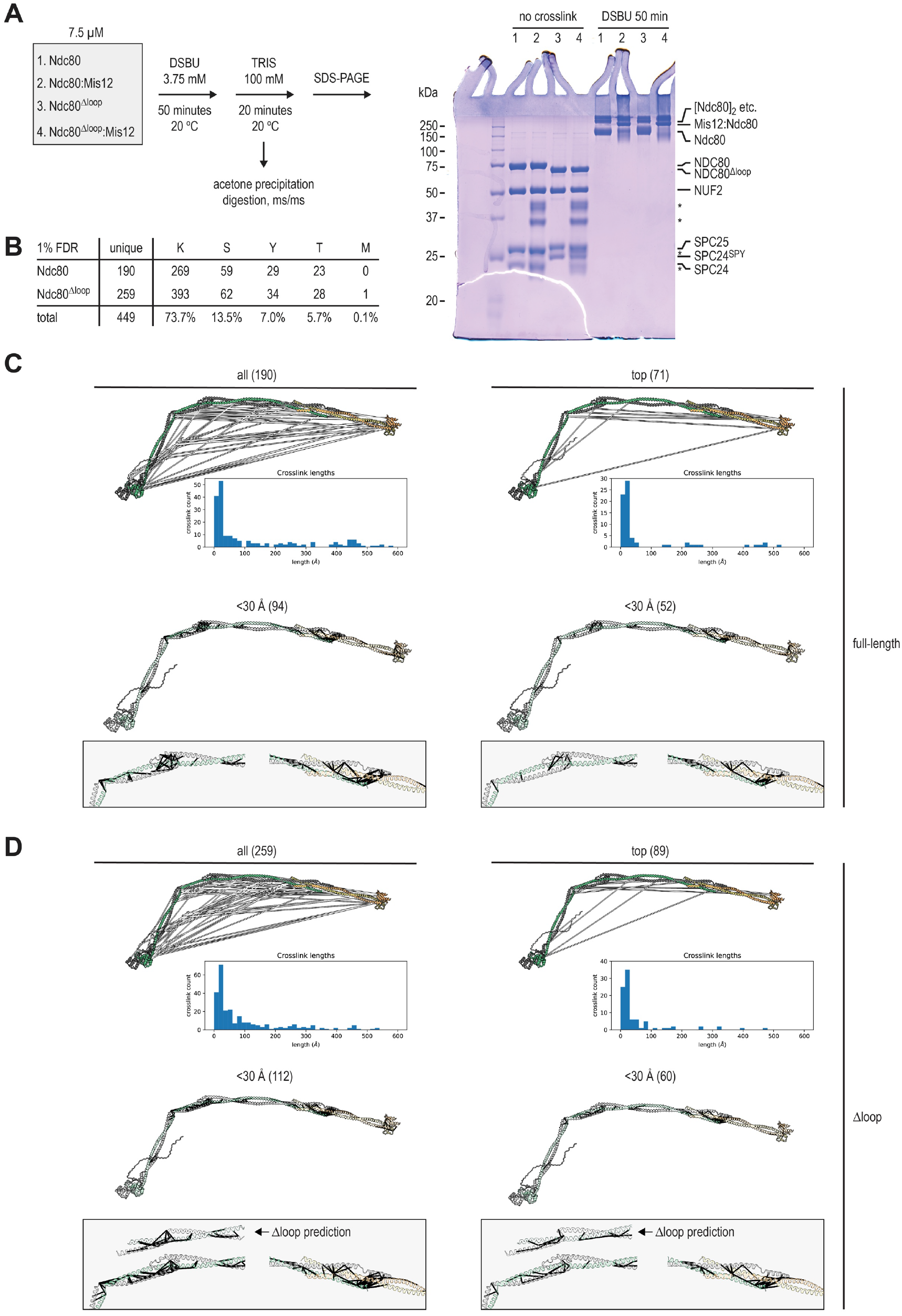
Chemical crosslinking followed by mass spectrometry and proximity maps. **A**) Crosslinking procedure and SDS-PAGE analysis of Ndc80, Mis12:Ndc80, Ndc80^Δloop^, and Mis12:Ndc80^Δloop^. The asterisks indicate the four subunits of the Mis12 complex. **B**) Analysis of side-chains crosslinked by DSBU in the various samples. M refers to the free NH_2_-terminus. **C**) Mapping of all (left) and top-scoring (right) crosslinks of full-length Ndc80 on the predicted structure of the full-length Ndc80 complex. A subset of crosslinks, all with a false-discovery rate below 1%, connect residues that are far apart in the extended Ndc80 structure. For instance, SPC25 K133 and K203 connect to various regions of the complex. Whether these long-distance crosslinks reflect transient compacted conformations of Ndc80 or transient inter-complex interactions is unclear. Lengths indicate C_α_-C_α_ distances. Crosslinks spanning a distance below 30 Å are shown separately, with magnifications of the loop and tetramerisation regions. **D**) Mapping of all (left) and top-scoring (right) crosslinks of Ndc80^Δloop^ on the predicted structure of the full-length Ndc80 complex. Crosslinks spanning a distance below 30 Å are shown separately, with magnifications of the loop and tetramerisation regions, as well as a prediction of the loopless Ndc80 region.

**Suppl. Figure 5.**
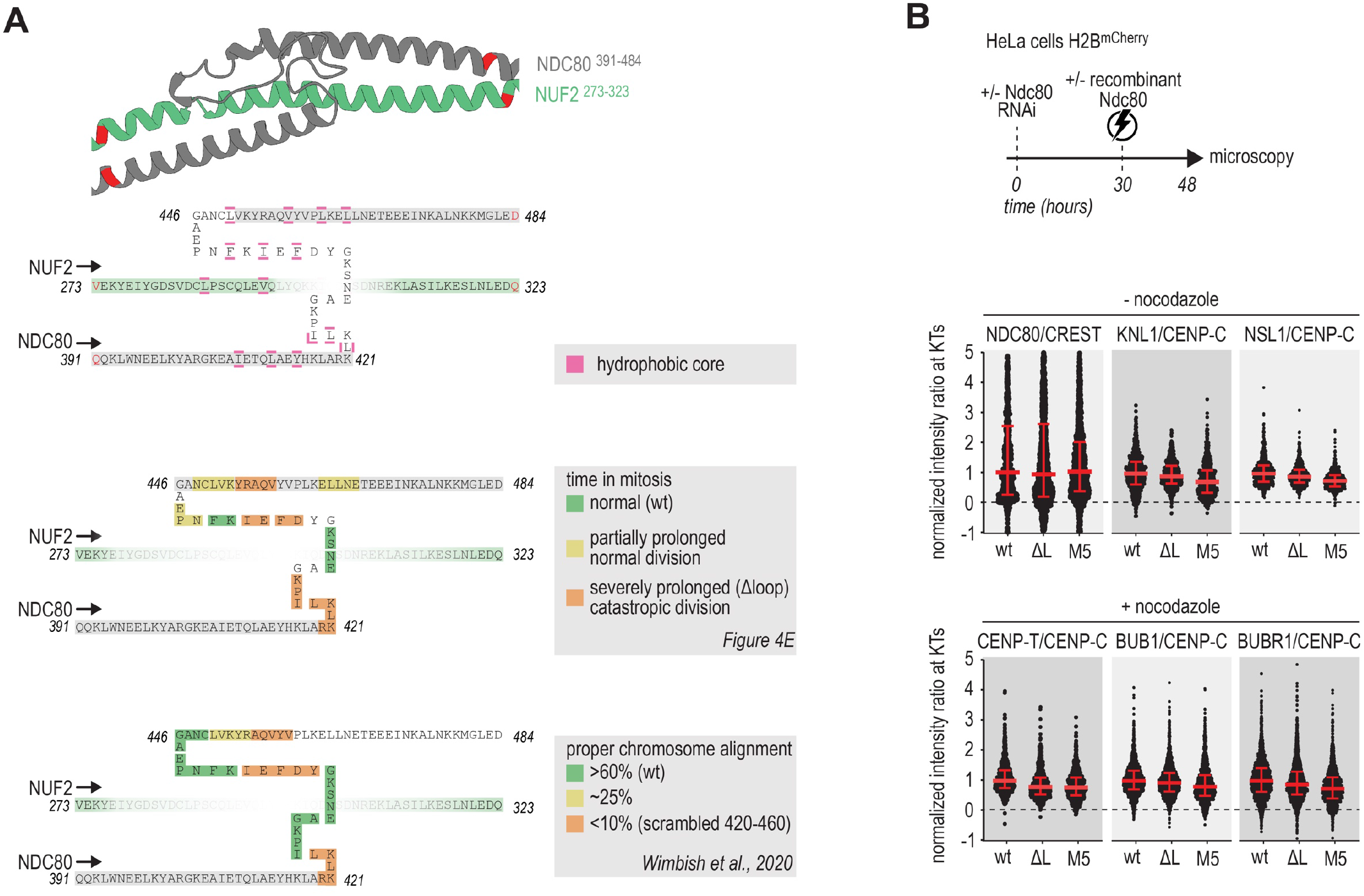
Electroporation efficiency and a comparison of loop mutants. **A**) Schematic representation of the predicted structure of the NDC80:NUF2 loop region with annotations to illustrate residues with side-chains that pack a hydrophobic core and mutants that interfere with chromosome congression as illustrated in **Figure 4E** and (Wimbish et al., 2020) **B**) Immunofluorescence quantification of various kinetochore and SAC proteins in cells that were treated as described in panel A. Nocodazole was added 15 hours after electroporation and 3 hours before fixation. Red lines indicate median value with interquartile range.

**Suppl. Figure 6.**
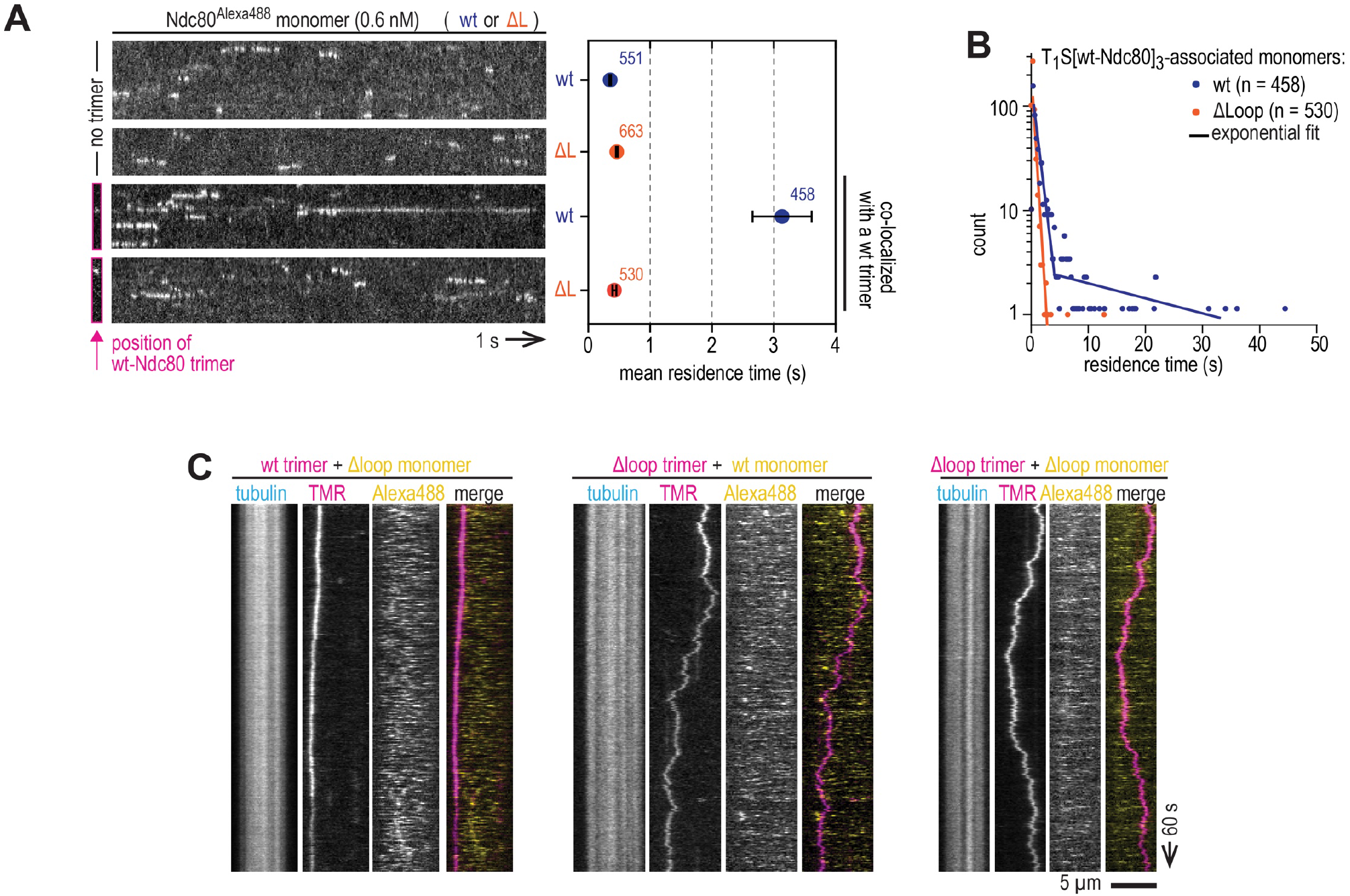
Loop-dependent binding between Ndc80 monomers and trimers on microtubules. **A**) Supplementary information for **Figure 5A-D**. High-speed recordings to quantify residence time of wild-type and loopless Ndc80 monomers. The top two kymographs show a microtubule with monomers binding and unbinding. The lower two kymographs show the binding and unbinding of Ndc80 monomers to a microtubule with a Ndc80 trimer. Since trimers are practically motionless on this timescale, only their initial location was recorded and indicated. Corresponding mean residence times and SEM. are shown. The number of analysed events is indicated. **B**) Distribution of residence times of wild-type and loopless Ndc80 monomers associating with microtubule-bound Ndc80 trimers. A single-exponential fit described the residence time of Ndc80^Δloop^, likely corresponding to Ndc80^Δloop^:microtubule off-rates. Residence time of full-length Ndc80 could be fitted with two exponents, likely corresponding to Ndc80:microtubule and Ndc80:trimer:microtubule off-rates. **C**) Typical kymographs showing Ndc80 trimers (magenta) and transiently binding Ndc80 monomers (yellow). Scale bars: vertical (100 s), horizontal (5 μm).

**Suppl. Figure 7.**
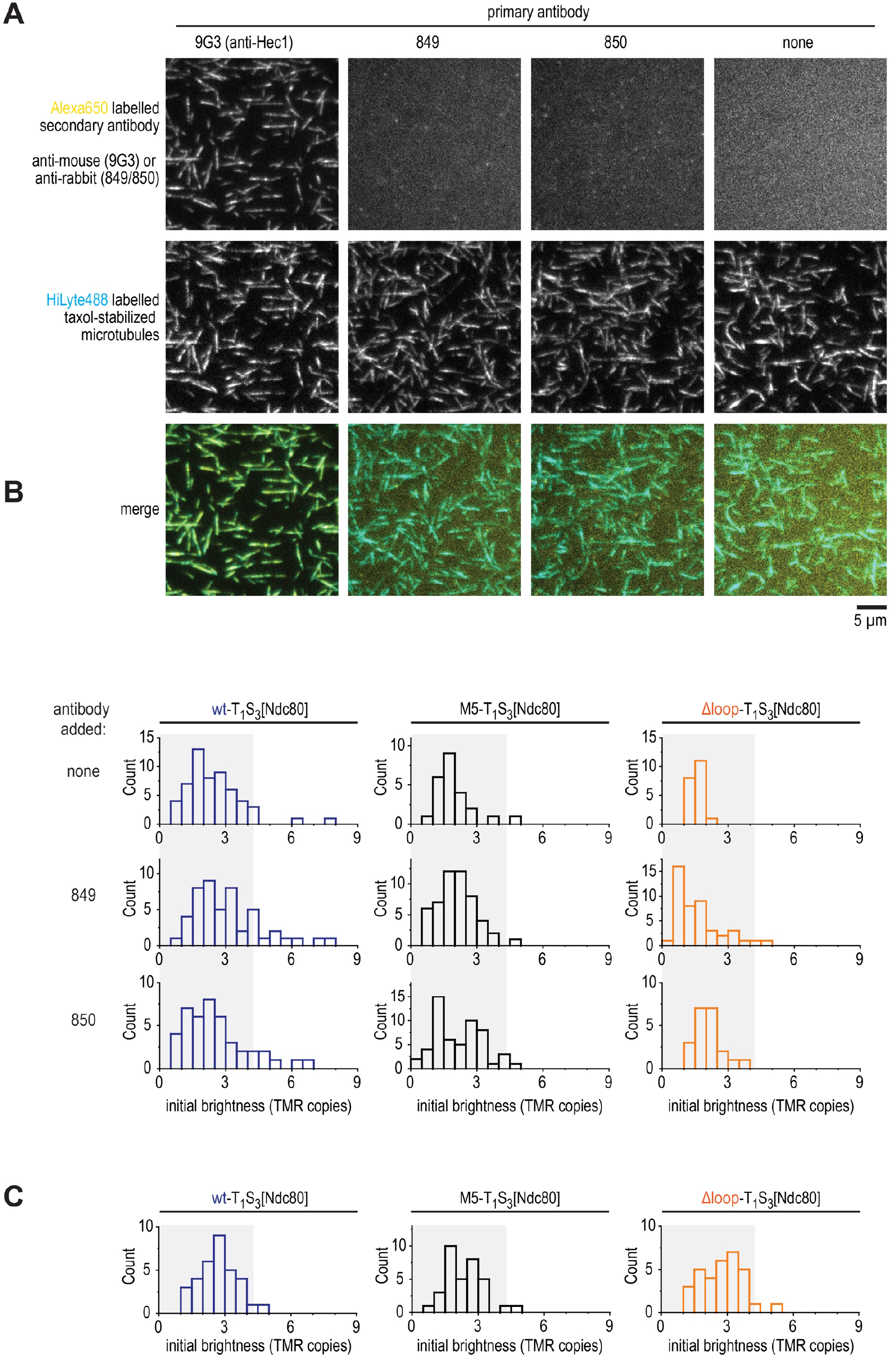
Characterisation of AB-849 and AB-850 *in vitro*. **A**) A fluorescently labelled secondary antibody was used to detect microtubule binding of primary antibodies in the absence of Ndc80. Scale bar: 5 μm. **B**) Distributions of initial brightness of Ndc80 trimers in absence and in presence of crosslinking antibodies. Shaded areas show data that was used to analyse diffusion. To enable experiments with antibodies, these experiments were performed without reducing agents. **C**) Comparable conditions as in panel A, but with the buffer including reducing reagents that was used for other *in vitro* experiments with microtubules (such as in **Figure 2D-E**).

**Suppl. Figure 8.**
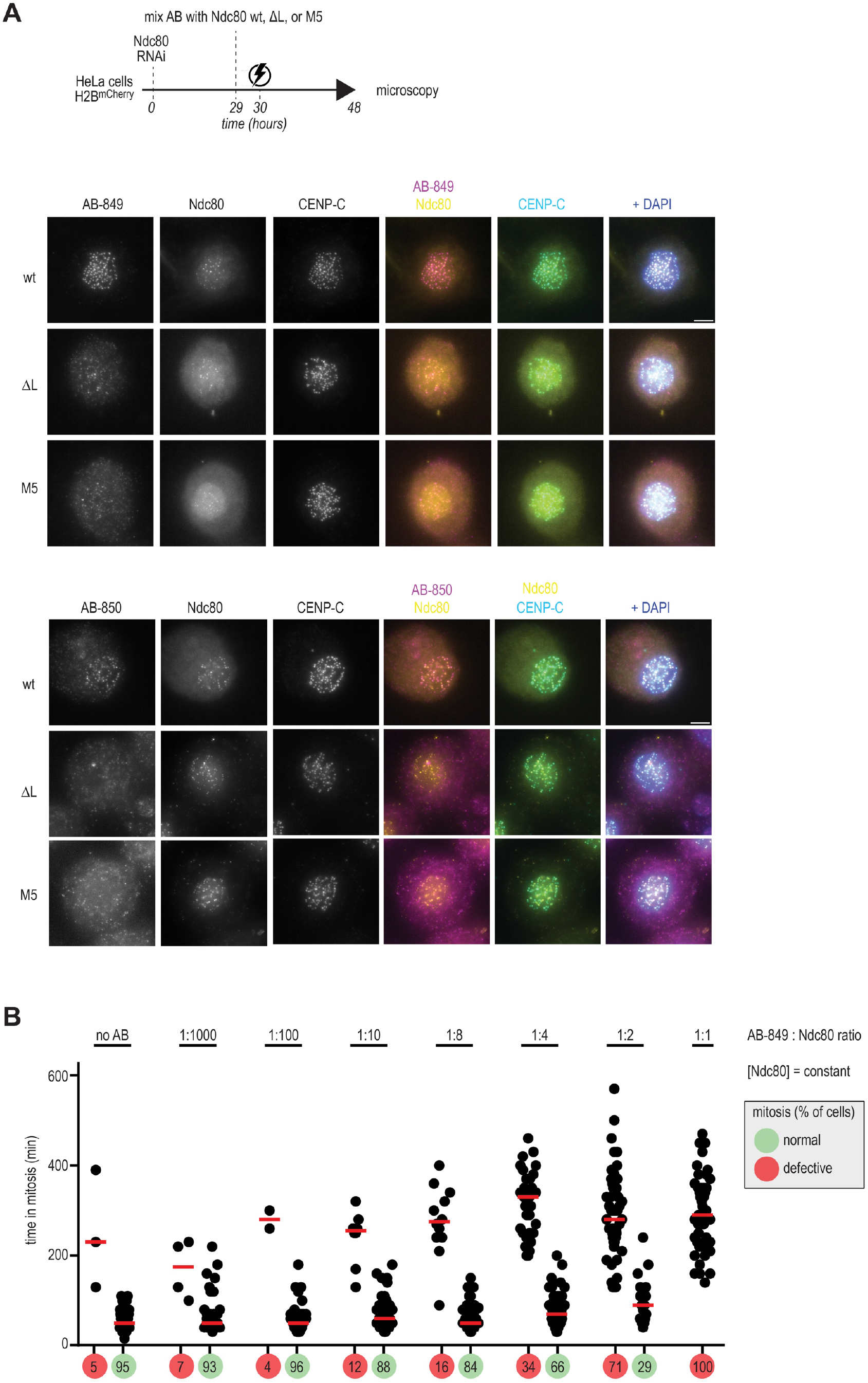
Characterisation of AB-849 and AB-850 *in vivo*. **A**) Experimental workflow as in **Figure 7B**. The localisation of AB-849, AB-850, NDC80, CENP-C, and DNA was analysed using immunofluorescence microscopy. Representative cells are shown. Scale bar – 5 μm. **B**) Experimental workflow as above, but with constant amounts of Ndc80 and decreasing amounts of AB-849 (the 1:1 ratio, approximately equimolar, was used in panel A and in **Figure 7B** with the Ndc80 complex). The time that cells spent in mitosis and the percentage of cells that divided normally (green) or divided abnormally after an arrest (red) are shown. Every dot represents a cell and red lines indicate median values.

